# Astrocyte networks and intercellular calcium propagation

**DOI:** 10.1101/248989

**Authors:** Jules Lallouette, Maurizio De Pittà, Hugues Berry

**Affiliations:** EPI BEAGLE, INRIA Rhône-Alpes, Villeurbanne, France

## Abstract

Astrocytes organize in complex networks through connections by gap junction channels that are regulated by extra‐ and intracellular signals. Calcium signals generated in individual cells, can propagate across these networks in the form of intercellular calcium waves, mediated by diffusion of second messengers molecules such as inositol 1,4,5-trisphosphate. The mechanisms underpinning the large variety of spatiotemporal patterns of propagation of astrocytic calcium waves however remain a matter of investigation. In the last decade, awareness has grown on the morphological diversity of astrocytes as well as their connections in networks, which seem dependent on the brain area, developmental stage, and the ultrastructure of the associated neuropile. It is speculated that this diversity underpins an equal functional variety but the current experimental techniques are limited in supporting this hypothesis because they do not allow to resolve the exact connectivity of astrocyte networks in the brain. With this aim we present a general framework to model intercellular calcium wave propagation in astrocyte networks and use it to specifically investigate how different network topologies could influence shape, frequency and propagation of these waves.

## 1 Introduction

An aspect of astrocytic Ca^2+^ signals is their ability to propagate as regenerative Ca^2+^ waves both intracellularly, i.e. within the same cell, and intercellularly, i.e. through different cells (Scemes and Giaume, 2006). In this fashion, processing of synaptic activity by Ca^2+^ in one region of an astrocyte can extend not only to other regions of the same cells but also to neighboring cells, potentially adding nonlocal interactions to the repertoire of neuron-glia interactions (De Pittà et al., 2012; Bazargani and Attwell, 2016).

Intercellular calcium waves (ICWs) have originally been reported in astrocyte cultures (Cornell-Bell et al., 1990; Blomstrand et al., 1999; Scemes et al., 2000) and then confirmed to also propagate in astrocytes in brain slices, (Sul et al., 2004; Schipke et al., 2002; Weissman et al., 2004) as well as in live rodents both in physiological (Kurth-Nelson et al., 2009; Kuga et al., 2011) and pathological conditions (Kuchibhotla et al., 2009). They can occur spontaneously (Nimmerjahn et al., 2004) or be evoked by exogenous stimuli (Ding et al., 2013; Sun et al., 2013) and either be restricted to few astrocytes (i.e. < 10 – 30) (Sul et al., 2004; Tian et al., 2006; Sasaki et al., 2011) or engulf hundreds of cells, while propagating in a regenerative fashion (Kuga et al., 2011). The reasons for this variety of modes of propagation remain however unknown. Besides differences in the experimental setups that include different brain regions, stimulus protocols or cellular Ca^2+^ responses, growing evidence suggests a further, previously unknown factor: the organization of astrocytes in variagated networks (Scemes and Giaume, 2006; Giaume et al., 2010).

Since the 1970s our understanding of intercellular communication between astrocytes has fundamentally changed from the notion that they are organized as a syncytium – a multinucleate mass of cytoplasm resulting from the fusion of cells – to the recognition that they are organized into networks with specific topology (Giaume et al., 2010). Neighboring astrocytes in different brain regions are indeed connected at their periphery by gap junctions (GJCs) – channels that allow the intercellular passage of ions and small molecules – and their anatomical domains only minimally overlap, as if they were tiling the brain space (Giaume and McCarthy, 1996; Bushong et al., 2002).

The mechanisms establishing whether two astrocytes are connected via GJCs are however nontrivial and far from being understood (Giaume, 2010). For example, the expression of connexins, in particular of Cx30 and Cx43 – the main proteins forming astrocytic GJCs (Giaume et al., 1991; Rouach et al., 2002; Koulakoff et al., 2008) –, is known to change across different brain regions (Blomstrand et al., 1999) and, in the case of Cx30 during development (Aberg et al., 1999; Montoro and Yuste, 2004). Even within the same brain region, GJC expression can considerably change across different structures. Indeed astrocytes within glomeruli of the olfactory bulb appear to be more connected than outside of these structures (Roux et al., 2011), and similar observations have been made in the somatosensory (barrel) cortex (Houades et al., 2008). While it is believed that this peculiar organization could define precise cellular and anatomical domains, neither the functional relevance of this specialized connectivity is known nor how it could ultimately affect astrocytic Ca^2+^ signaling (Pannasch and Rouach, 2013).

Current experimental techniques do not allow to resolve the exact connectivity (topology) of astrocyte networks in the brain and thus are not helpful to address these aspects. In this perspective, computational approaches can provide a valuable tool to investigate general topological principles underpinning ICW propagation (or lack thereof) in astrocyte networks. Here we review some of these approaches in the context of 2-dimensional and 3-dimensional astrocyte networks, leveraging our modeling arguments on observations from dedicated experiments in mixed neuron-glia cultures.

## 2 Astrocyte network modeling

### 2.1 General framework

Modeling of astrocyte networks may be pursued in different ways depending on what extent we want to take into account astrocyte anatomy. Astrocytes have indeed complex anatomy, with multiple primary processes irradiating from their somata and branching into secondary and tertiary processes that end in a myriad of tiny lamellipodia and filopodia (Theodosis et al., 2008). Accordingly, ICWs can be described as continuous waves that gradually propagate through this complex medium ensuing from this intricate network of astrocytic processes. While the mathematical theory of these waves is well developed (Falcke, 2004), this approach is however limited by the lack of tools to resolve the fine structure of astrocytic secondary and tertiary processes, except for simple setups of cell cultures (Kang and Othmer, 2009).

Alternatively, we may consider only somatic activation and describe ICWs as propagating waves that hop from one astrocyte to neighboring ones in a coarse-grained fashion, that is counting the number of cells that are activated by a wave rather than the spatial extent to which the wave propagates through the intricate ensemble of astrocytic processes. In this fashion, astrocyte somata are the nodes of the network, whose activation can be described in principle by the (time) evolution of two state vectors: **a** = (*C*,…) which lumps the astrocyte’s intracellular Ca^2+^ concentration, (*C*), along with possible other variables that control it, like gating variables of intracellular channels that regulate Ca^2+^ release from the endoplasmic reticulum, or Ca^2+^ buffers that prevent Ca^2+^ accumulation in the cytosol (Falcke, 2004, p. 291); and s which accounts for Ca^2+^-mobilizing signals that are responsible for regenerative propagation of ICWs.

Denoting by 𝒩_*i*_ the set of astrocytes that are neighbors of cell *i* in the network, the equations of these state vectors associated with cell *i* generally read

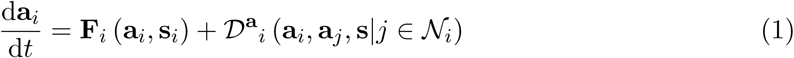

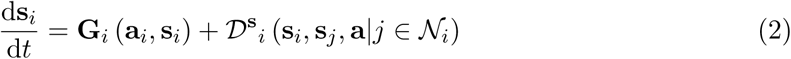

where the vector functions **F**_*i*_ and **G**_*i*_ are typically nonlinear, and the terms 𝒟^**a**^_*i*_ and 𝒟^s^_*i*_ account for exchange of chemical species (lumped in **a**_*i*_ and **s**_*i*_) between the *i*-th astrocyte and its neighbors.

While equations 1 and 2 do not account for the totality of models of ICWs, they can nevertheless describe a large class of whole cell models often used in the study of astrocytic ICWs, some example of which we also consider in this chapter. In particular, depending on the choice of the astrocyte model, besides cytosolic Ca^2+^, the components of the state vector a and the vector function **F** can include the Ca^2+^ concentration in the endoplasmic reticulum (Dupont and Goldbeter, 1993), the state variables of the Ca^2+^ release channels and their dynamics (De Young and Keizer, 1992; Li and Rinzel, 1994; Tang and Othmer, 1994; Sneyd et al., 1998; Höfer et al., 2002; Stamatakis and Mantzaris, 2006). Similarly, in addition to the proper Ca^2+^ mobilizing second messenger molecules, the state vector s and the vector function **G** may also include the state variables for the kinetics of the receptors that control the generation of those second messengers (Kummer et al., 2000; Höfer et al., 2002; Stamatakis and Mantzaris, 2006; Ullah et al., 2006a) as well as for other molecular signals involved in the intracellular regulation of such messengers (Chay et al., 1995; Bennett et al., 2005). In many situations, dynamics of the components of **a** and **s** are interdependent as mirrored by the fact that **F** and **G** in the above equations are functions of both state vectors. This is obvious for second messengers that control intracellular Ca^2+^ dynamics, but it is also often the case that Ca^2+^ itself can regulate multiple aspects of the dynamics of those second messengers (Chay et al., 1995; Höfer et al., 2002; Ullah et al., 2006a). This may also be the case for the two exchange terms 𝒟^**a**^ and 𝒟^**s**^, which generally account for intra‐ and inter-cellular diffusion of Ca^2+^ along with Ca^2+^-mobilizing second messenger molecules (Höfer et al., 2002; Stamatakis and Mantzaris, 2006; Edwards and Gibson, 2010), insofar as the rate of such diffusion may depend on these latter, for example through Ca^2+^-dependent buffers (Kupferman et al., 1997; Sherman et al., 2001) or secondary reactions involving second messenger molecules (Dupont and Erneux, 1997; Stamatakis and Mantzaris, 2006).

In general, two are the routes for chemical exchange between astrocytes that are involved in ICWs: one is by intracellular diffusion of Ca^2+^ and the second messenger molecule inositol 1,4,5-trisphosphate (IP_3_) through GJCs, the other one is by Ca^2+^-dependent ATP release from astrocytes into the extracellular space (Scemes and Giaume, 2006). Both routes, although brought forth by different biochemical reactions, promote IP_3_-triggered Ca^2+^-induced Ca^2+^ release (CICR) from the endoplasmic reticulum, which is the main mechanism of Ca^2+^ signaling in ICWs (Nedergaard et al., 2003). This is obvious in the intracellular route whereby IP_3_ is supplied to resting cells via GJCs. In the extracellular route instead, this is mediated by the activation of metabotropic purinergic receptors which, akin to glutamatergic receptors (Chapter 5), trigger IP_3_ production (and CICR) by G_q_ protein-mediated hydrolysis of phosphoinositol 4,5-bisphosphate (Scemes and Giaume, 2006).

From a modeling perspective, the fact that CICR is the main mechanism of Ca^2+^ signaling in ICWs, allows to replace equation 1 by any model of CICR (Chapter 2). Moreover it is also possible to neglect intracellular Ca^2+^ diffusion because free Ca^2+^ is rapidly buffered in the astrocyte cytosol, thereby minimally leaking through GJCs (Allbritton et al., 1992; Sneyd et al., 1998; Höfer et al., 2002). This allows to simplify equation 1 by setting 𝒟^**a**^_*i*_ = 0, and only leaves to specify **s**_*i*_ and equation 2. With this regard, both intracellular IP_3_ and extracellular ATP can contribute together to ICW, with their relative involvement likely depending on regional, developmental and experimental conditions (Scemes and Giaume, 2006). Nonetheless, because hereafter we aim to characterize how different connections between astrocytes could affect ICW propagation, we limit our analysis to the consideration of GJC-mediated IP_3_ diffusion only. The reader who is interested in modeling purinergically mediated ICWs may refer to Bennett et al. (2005) and MacDonald et al. (2008) for astrocyte network models that consider ATP signaling only, or alternatively to Iacobas et al. (2006); Stamatakis and Mantzaris (2006); Kang and Othmer (2009) and Edwards and Gibson (2010) for models that include extracelluar ATP signaling in combination with intracellular IP_3_ diffusion.

### 2.2 Gap junction-mediated IP_3_ diffusion

In its most general form, the flux of IP_3_ (I) mediated by diffusion from one astrocyte *i* to a neighboring one *j* (*J*_*ij*_) can be thought as some function *ϕ* of the IP_3_ gradient between the two cells, i.e. Δ_*ij*_*I* = *I*_*i*_ – *I*_*j*_ (Crank, 1980), so that

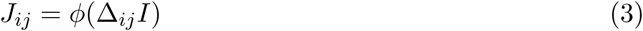

In the simplest scenario of short distance and/or fast diffusion, the intracellular environment along the pathway from cell *i* to *j* may be assumed homogeneous so that *ϕ* is linear (Sneyd et al., 1994; Falcke, 2004), and *J*_*ij*_ is accordingly described by Fick’s first diffusion law, i.e.

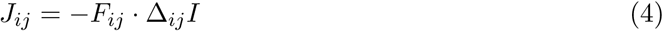

where *F*_*ij*_ is the diffusion coefficient. In practice however, IP_3_ diffusion between astrocyte somata could be more complicated. This is because connections between astrocytes through GJCs are mostly at the cell distal processes (Giaume et al., 2010) whose complex morphology and narrow intracellular space (Witcher et al., 2007; Pivneva et al., 2008) could considerably hinder IP_3_ diffusion from/to somata. Moreover, GJCs cluster at discrete sites of these processes (Nagy and Rash, 2000), thereby constraining the diffusion pathway of IP_3_ from one cell to another. Finally, IP_3_ production and degradation in the processes could either promote IP_3_ transfer between cells or hamper it. In this fashion, the ensemble of astrocytic processes and GJCs interposed between cell somata could equivalently be regarded as a diffusion barrier for IP_3_ exchange between cells, and accordingly, IP_3_ diffusion between cells could be inherently nonlinear. This scenario is further substantiated by growing experimental evidence suggesting that GJC permeability could be actively modulated by various factors, including different second messengers (Harris, 2001). With this regard, the permeability of Cx43, a predominant connexin in astrocytic GJCs (Nagy and Rash, 2000), could be modulated for example by phosphorylation by protein kinase C (Bao et al., 2004; Sirnes et al., 2009; Huang et al., 2013). Because the same kinase also takes part in IP_3_ degradation as well as in Ca^2+^ signalling (Codazzi et al., 2001; Irvine and Schell, 2001), this possibility ultimately hints that GJC permeability could also depend on IP_3_ signaling, whose dynamics is notoriously nonlinear (Chapter 5).

The above arguments support the choice of a nonlinear *ϕ* in equation 3. With this regard then, we may assume that IP_3_ diffusion between two astrocytes, *i* and *j*, is a threshold function of the IP_3_ gradient between somata of those cells, whose strength is bounded by the maximal GJC permeability. In this way, a possible expression for *J*_*ij*_ is (Goldberg et al., 2010):

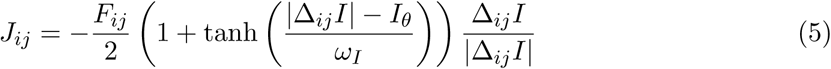

where *I*_*θ*_ represents the threshold gradient for which effective IP_3_ diffusion occurs; whereas *ω*_*I*_ scales how fast *J*_*ij*_ increases (decreases) with Δ_*ij*_*I* (see Figure 1C). The parameter *F*_*ij*_, which in the linear approximation sets the slope of *J*_*ij*_ (equation 4), here fixes instead the magnitude of the maximum diffusion flux.

**Figure 1:**
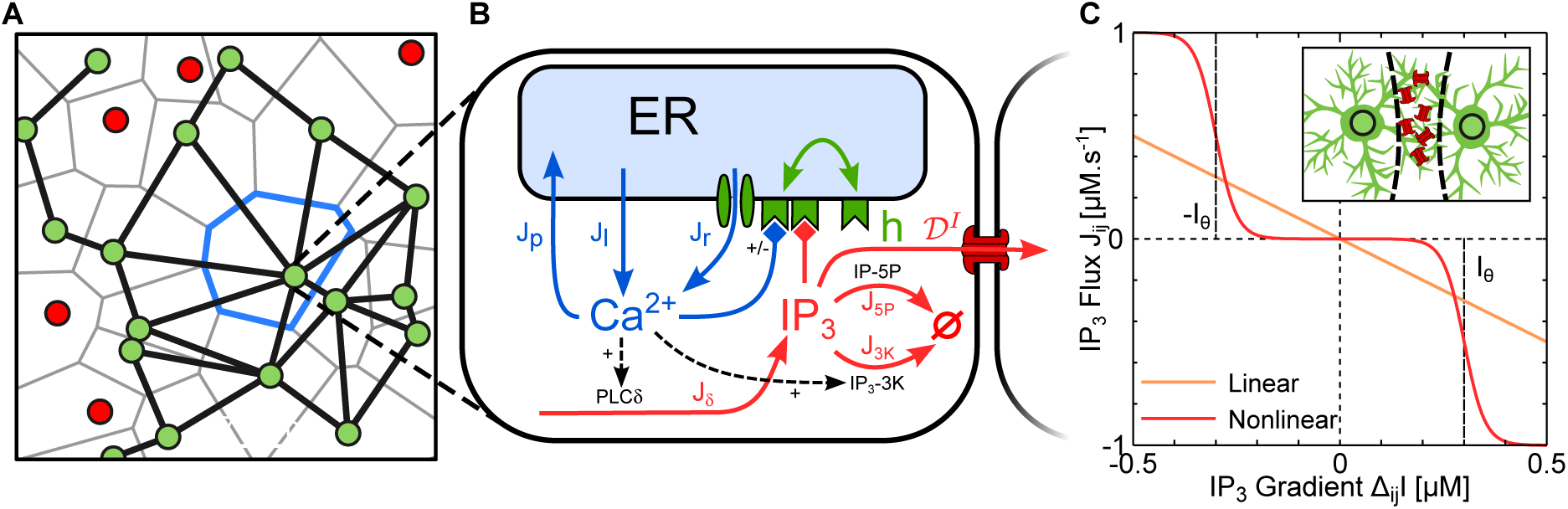
Modeling astrocyte networks. **A** Double immunostaining of a cultured neuron-glial network by the neuronal marker NeuN (*red*) and the astrocytic marker GFAP (*green*). Astrocyte anatomical domains (like the one delimited in *blue*) are reconstructed by Voronoi tessellation (*gray lines*) of the network, only considering neurons and astrocytes activated by electrical stimulation (see Wallach et al., 2014, for details). Neighboring astrocytes are assumed to be connected by GJCs when their Voronoi cells are contiguous. **B** Schematic representation of the biophysical network model. Individual astrocytes are described by the well known ChI model for astrocytic Ca^2+^-induced Ca^2+^ release from the endoplasmic reticulum (De Pittà et al., 2009), while their coupling with neighboring cells is by intercellular IP_3_ diffusion by GJCs (𝒟^I^). In some simulations, we also consider glutamate-mediated IP_3_ production by PLCβ to account for synaptically-evoked Ca^2+^ signals (not shown). **C** Differently from classic (linear) diffusion (yellow line), IP_3_ diffusion between astrocytic somata is modeled by a nonlinear (sigmoid) function of IP_3_ gradient between cells. (*Inset*) This choice takes into account that most GJCs (in *red*) are in the processes of astrocytes at the border of their anatomical region.

### 2.3 Network topology

We have introduced so far a general framework to model individual astrocytes (as nodes) of the network (equations 1 and 2), and their connections by GJC-mediated exchange of IP_3_ (equation 5). In order to complete our description of the astrocyte network we must then specify the connections of each cell with others in the network.

Generally speaking, astrocytes networks can develop in one, two or three dimensions. The simplest scenario of 1d networks, that is astrocyte chains, is useful to investigate how cellular properties could affect ICWs. With this regard for example, cellular mechanisms controlling CICR rate (Höfer et al., 2001; Ullah et al., 2006b), the type of encoding by Ca^2+^ oscillations (Goldberg et al., 2010) or GJC permeability (Matrosov and Kazantsev, 2011) have been shown to critically control the number of astrocytes recruited by ICWs. These results have also been confirmed by 2d astrocyte network models, that are a valuable tool to investigate the rich variety of patterns of propagation of astrocytic ICWs observed in cell cultures (Sneyd et al., 1994, 1995a,b; Sneyd and Sherratt, 1997; Sneyd et al., 1998; Höfer et al., 2002; Shuai and Jung, 2003; Dokukina et al., 2008). The vast majority of these models however assumes a simplified arrangement of astrocytes on a regular lattice focusing on the CICR nonlinearity to exploit complex modes of ICW propagation. Only few studies have explored instead the potential role of network topology on ICW nucleation and propagation. Dokukina et al. (2008) considered for example small ensembles of three or four interconnected astrocytes, showing that only some connection schemes, among all possible ones, can favor ICW generation, while variations in GJC permeability can hamper ICW propagation regardless. More recently Wallach et al. (2014) and Lallouette et al. (2014) attempted to extend this analysis to large 2d and 3d networks with the aim to derive principles of astrocyte ICW propagation driven by network topological features. The main results of these two studies are reproduced in the next section to illustrate our modeling approach.

## 3 Biophysical modeling of intercellular Ca^2+^ waves

### 3.1 Ca^2+^ signaling in mixed neuronal and astrocytic cultures

As a first example of application of our modeling approach introduced in the previous section, let us consider the task of modeling Ca^2+^ signaling in cultured mixed neuronal and astrocytic networks. With this regard, we consider the experiments by Wallach et al. (2014) where this common experimental setup was used in combination with electrical stimulation of neural activity, at rates from 0.2 to 70 Hz, to trigger astrocytic Ca^2+^ signaling. In those experiments, bath perfusion with antagonists of astrocytic glutamate receptors was shown to block evoked Ca^2+^ activity in astrocytes, and so it was concluded that this signaling ensued from synaptically-released glutamate, which likely activated astrocytic type I metabotropic glutamate receptors. Based on these considerations, we replace **a, s, F** and **G** in equations 1 and 2 by the biophysical model of glutamate-mediated astrocytic Ca^2+^ signaling discussed in Chapter 5 (see also De Pittà et al., 2009), so that the generic *i*-th astrocyte in the cultured network is described by (Figure 1B):

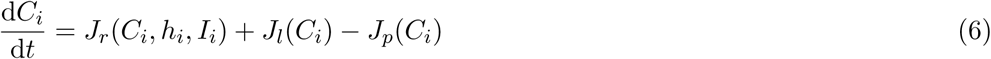

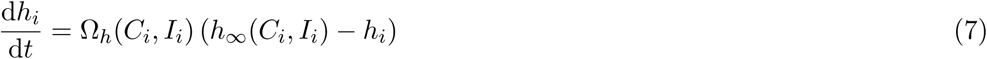

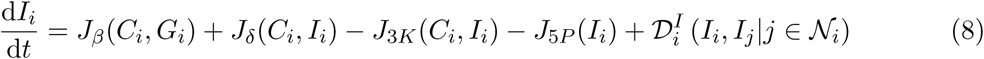

where intercellular exchange of IP_3_ in the last equation is taken equal to the sum of individual diffusion fluxes (equation 5) between astrocyte *i* and its connected neighbors, i.e.

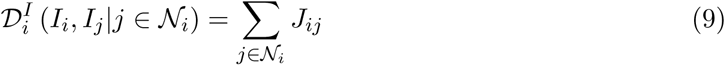

To complete the model, we assume that glutamate concentration (*G*_*i*_) in the medium surrounding astrocytic receptors instantaneously increases for each electrical pulse delivered at time *t*_*k*_, proportionally to the available synaptic glutamate resources (*g*), and exponentially decays between pulses at rate Ω_*G*_, mimicking glutamate clearance by diffusion in the extracellular space (Clements et al., 1992), i.e. *G*_*i*_(*t*) ≈ Σ_*k*_ *g*(*t*_*k*_) exp (–Ω(*t* – _*k*_)) Θ (*t* – *t*_*k*_) where Θ(·) is the Heaviside function (Wallach et al., 2014).

To build realistic astrocytic networks, we then borrow the argument that astrocytes likely tile the space of neuronal networks they are in by their non-overlapping anatomical domains (Bushong et al., 2002). In this fashion, adjacent astrocytes are more likely to be connected by GJCs than cells that are far apart (Giaume et al., 2010). Accordingly, we consider immunostaining images of the cultured networks, like the one in Figure 1A where somata of neurons and astrocytes are respectively marked by *red* and *green circles*, and construct the Voronoi diagram (*gray lines*) associated with every cell in the network. This diagram partitions the network into as many regions as the cells taken into account, where each region may be regarded as an estimate of the anatomical domain (*blue lines*) of the cell that it contains (Wallach et al., 2014; Galea et al., 2015; Sánchez-Gutiérrez et al., 2016). Thus considering only the regions associated with activated astrocytes, in our modeling we assume neighboring astrocytes to be connected by GJCs only if their corresponding Voronoi regions share a border. We repeat this procedure for all cell cultures imaged by Wallach et al. (2014) and consider Ca^2+^ signals evoked by repetitive neural (synaptic) stimulation of our model astrocytes. Since we are interested in the possible influences of different connections between astrocytes on their Ca^2+^ response, we model all astrocytes in each culture identically, varying only their connectivity according to their Voronoi tessellation.

Time-frequency characterization of astrocytic Ca^2+^ responses recorded in experiments by Wallach et al. (2014) are shown in Figure 2A, where two classes of responses may be recognized. *Type I* responses (*top row*) are characterized by astrocyte activation at relatively high frequency of neural activation (*top bars*), while the frequency of Ca^2+^ oscillations does not significantly change as the rate of neural stimulation increases. *Type II* responses instead, can be observed for slightly lower frequencies of neural activation (*bottom row*), but are distinguished by Ca^2+^ oscillations whose frequency increases with neural stimulation, reaching values that are generally higher than in type I responses.

**Figure 2:**
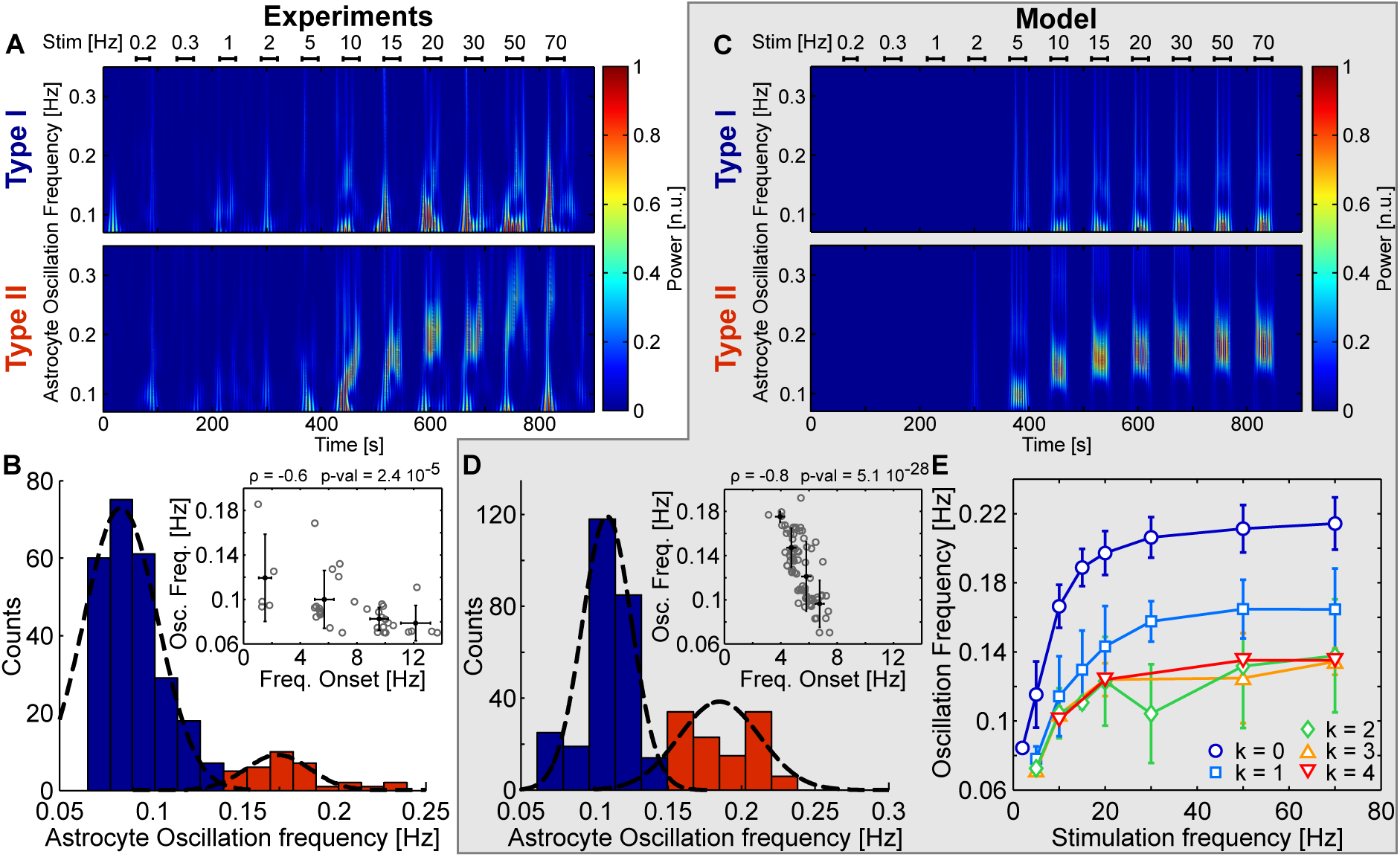
Regulation of Ca^2+^ oscillations in individual astrocytes by GJCs. **A** Time-frequency analysis of Ca^2+^ oscillations evoked in individual astrocytes in cell cultures by repetitive electrical stimulation of neurons at different rates (“Stim.”, *top bars*). Some astrocytes display low frequency oscillations (Type I responses, *top row*), while others show higher oscillation frequencies that increase with the rate of neural stimulation (Type II responses, *bottom row*). **B** Distribution of maximal frequencies of recorded astrocytic Ca^2+^ oscillations. Type I oscillations (*blue bars*) and Type II responses (*red* bars) are neatly separated and fitted by Gaussian distributions with different mean and variance (*black dashed curves*). (Inset) The frequency of Ca^2+^ oscillations is negatively correlated with the threshold rate of neural stimulation (“Onset freq.”) required to trigger them, so that high frequency/Type II responses are generally observed at lower onset rates of neural activation than low frequency/Type I responses. *Grey circles* denote single astrocytic responses; *black dots* represent means for representative onset rates; error bars denote standard deviation. **C, D** Results from numerical simulations of astrocytic networks reconstructed by immunostaining images (see Figure 1). *E* Maximum frequency of simulated Ca^2+^ oscillations as a function of the rate of neural stimulation for astrocytes with different number of connections with unstimulated neighbors 〈*k*〉. Increasing the stimulus rate increases the frequency of Ca^2+^ oscillations which plateaus for high rates of stimulation, independently of *k*. The height of this plateau however strongly depends on the number of connections between astrocytes so that unconnected astrocytes (*k* = 0) tend to oscillate much faster than connected ones (*k* > 0). Data points±errorbars denote mean values±standard deviation for *n* = 130 simulated astrocytes. Adapted from Wallach et al. (2014).

Consideration of the distribution of the maximum frequency of Ca^2+^ oscillations of all recorded responses in Figure 2B shows that roughly 80% of recorded astrocytes exhibited responses of type I (*blue bars*), with Ca^2+^ oscillating on average at most at ~ 0.1 Hz (*left peak* of the *dashed curve*); while the remaining astrocytes displayed type II Ca^2+^ responses (*red bars*), with approximately doubled average maximum frequency, i.e. ~ 0.2 Hz (*right peak* of the *dashed curve*). In parallel it may be appreciated from the *inset* how this maximum frequency of Ca^2+^ oscillations inversely correlates with the rate of neural stimulation, with high frequency/type II-like Ca^2+^ oscillations triggered by lower rates of neural stimulation than low frequency/type I-like oscillations.

To check consistency of our modeling approach, we reproduce in Figures 2C and 2D the previous results, yet based on Ca^2+^ responses generated by numerical simulations of our network models built by the procedure above described. It may be appreciated how, despite some inevitable quantitative differences, these figures qualitatively reproduce the essential features of experimental observations on the two types of astrocytic Ca^2+^ responses presented in Figures 2A and 2B, and thereby prove the effectiveness of our approach in modeling Ca^2+^ signaling in cultured neuron-glial networks.

To characterize the effect of astrocytic connectivity on individual Ca^2+^ responses, we next consider the maximum frequency of astrocytic Ca^2+^ oscillations simulated for different rates of neuronal stimulation, distinguishing among responses based on the number of connections (*k*) of individual astrocytes to unstimulated neighbors – the reason of this specific choice of neighbors will be clarified in the following sections. The results of this analysis are reported in Figure 2E where three observation may be made. First, it may be appreciated how unconnected astrocytes (*k* = 0, *dark blue curve*) display the highest oscillation frequency for all rates of neural stimulation. On the contrary, as the number of connections to unstimulated neighbors increases, the maximum frequency of oscillations decreases. Second, the threshold rate of neural stimulation to trigger astrocyte Ca^2+^ activation, tends to increase with the number of connections, being as low as ~ 2 Hz for unconnected astrocytes (*leftmost blue circle*) while increasing up to ~ 10 Hz for cells with *k* = 4 unstimulated connected neighbors (*leftmost downward red triangle*). Finally, the shape of the curves for different k values changes. For those astrocytes characterized by *k* ≤ 3 the maximum frequency of Ca^2+^ oscillations nonlinearly increases with the rate of neuronal stimulation, reaching values as high as ~ 0.2 Hz in unconnected (*k* = 0) cells. But this increase is progressively reduced as k grows larger, till it becomes almost negligible as in the case of astrocytes with *k* = 4 (*red curve*), for which the maximum frequency of Ca^2+^ oscillations is ~ 0.1 Hz, independently of the rate of neural activity.

Combining these considerations with the experimental results in Figures 2A and 2B, it is striking to correlate unconnected astrocytes, or astrocytes with a low number of unstimulated connected neighbors, with low onset rate/high oscillation frequency/type II responses; and vice versa, astrocytes with a high number unstimulated connected neighbors with high onset rate/low oscillation frequency/type I responses. At the lower extremum of this spectrum of astrocytic connectivity, we find unconnected astrocytes, which represent a minority, up to ~ 20% of astrocytes in cell cultures (Rouach et al., 2000), to likely account for the 0.2 Hz peak in the distribution of Ca^2+^ oscillations in Figure 2B. Conversely, at the higher extremum of the spectrum we find those astrocytes with *k* = 4, insofar as they could mainly account, together with some astrocytes with 2 ≤ *k* < 4, for the 0.1 Hz peak of that distribution.

To summarize, our hitherto analysis hints that the way astrocytes are connected can affect how they respond to neural activity, controlling the threshold neural stimulation required for their activation and the frequency of ensuing Ca^2+^ oscillations (Wallach et al., 2014). These results have been obtained however in a somewhat simplified setup which is that of 2d-like cultured astrocyte networks. In practice astrocytes are organized in three-dimensional networks in the brain, thus we following extend our biophysical modeling approach to address how topological differences in 3d networks could ultimately influence astrocytic Ca^2+^ activity.

### 3.2 Ca^2+^ wave propagation in 3d astrocyte networks

To model realistic 3d astrocyte networks we need to specify not only the topology of these networks but also, preliminarily, the arrangement of all cells in the 3d physical space. For 2d-like networks, such as the cultured networks modeled in the previous section, this task is eased by the possibility to exactly map every cell position, for example by immunostaining and post-fixation optical microscopy of the network. However, this information is currently not accessible experimentally for 3d astrocytic networks, although recent advances of connectomics could soon fill in this gap (Kasthuri et al., 2015). On the other hand, astrocyte arrangement in some brain regions has recently been characterized on statistical bases, and we following base our modeling on these data as originally described by Lallouette et al. (2014).

We consider a pool of *N* = 11^3^ astrocytes modeled by equations 6-8 and position them on a cubic lattice with internode distance *a*. We then jitter each cell location in the lattice by Gaussian noise with zero mean and variance *σ*^2^. In doing so, we choose *a* and *σ*^2^ to minimize the squared error with respect to experimental values of mean (50 μm), minimum (20μm), and coefficient of variation of cell distance (~ 0.25) (Sasaki et al., 2011, see Table C2 for specific parameter values). After positioning astrocytes in the physical space, we specify their connections, considering different topologies (Figure 3A), ranging from (i) strongly spatially-constrained networks such as *link radius networks*, where an astrocyte connects to all cells located within a given distance from its center, to (ii) completely spatially-unconstrained, random networks of *Erdős-Rényi topology*. In between these extrema we also consider (iii) *regular degree networks* where each astrocyte connects to its *k* nearest-neighbors, where *k* is the network *degree k;* (iv) *shortcut networks* are obtained by rewiring a fraction of the connections (chosen at random) of a regular degree network; replacing the destination cell of the original connections by a randomly-chosen cell of the network (independently of the distance); and (v) *spatial scale-free networks* where astrocyte degree follows a power-law distribution dependent on cell degree and distance (see Appendix A).

**Figure 3:**
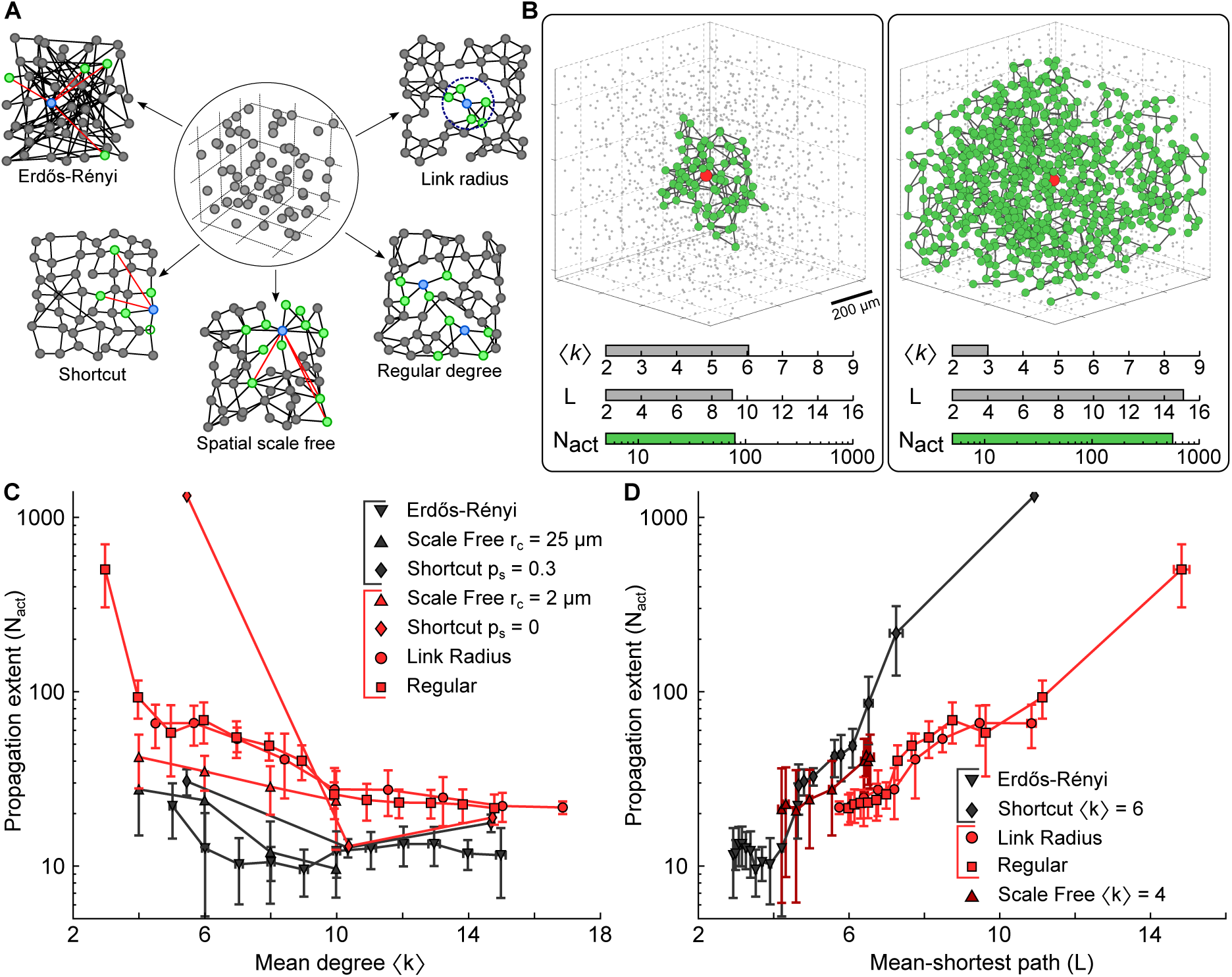
ICW propagation in 3d astrocyte networks. **A** Modeling procedure. (*Middle panel*) Astrocytes (*circles*) are first positioned on a cubic lattice, and their positions are randomly jittered to match experimentally-derived statistics of intercellular distances. (*Other panels*) Then they are connected by GJCs (*black lines*) according to different topologies (shown in 2d for clarity). *Red links* denote long-distance connections; cells in *green* are connected neighbors of *blue* cells. **B** Simulations of ICWs in two regular networks with different mean degree (〈*k*〉) and mean shortest path length (*L*). Both networks contained the same number of identical cells (*N* = 11^3^), and ICW stimulation was delivered to the central *red* cell in the same fashion (Appendix A), so that the 10-fold difference in the number of cells activated by the two ICWs (*N*_*act*_, *green circles*) only depended on different connections between the two networks. **C** Extent of ICW propagation as a function of the network’s mean degree 〈*k*〉 and **D** mean shortest path **L**. In both cases, spatially-constrained topologies (*red markers* e.g. link radius, regular degree and shortcut with *p*_*s*_ = 0 or 〈*k*〉 = 4) can support ICW propagation up to several hundreds of astrocytes, whereas spatially-unconstrained topologies cannot (*black markers* e.g. Erdős-Rényi, shortcut with *p*_*s*_ > 0, scale-free). Scale-free networks in **D** (*dark red upward triangles*) can either be spatially constrained or unconstrained, depending on the value of *r*_*c*_. Data points±errorbars correspond to mean values±standard deviation over *n* = 20 different realization of the network for fixed statistical parameters. Adapted from Lallouette et al. (2014). Model parameters as in Table C1.

Let us now consider the propagation of ICWs in the model networks and study how the extent of this propagation, quantified by the number of astrocytes activated at least once by an ICW (*N*_*act*_), depends on network topology. With this aim, we trigger ICW propagation in our model networks stimulating CICR in the cell in the center of the 3d space of the network to minimize boundary effects (see Appendix A for details). Two examples of ICWs triggered in this fashion are shown in Figure 3B for two different regular degree networks. The difference in the number of activated cells, represented by *green* circles, is striking and suggests that simple variations in network topology could account for large variability in ICW propagation. In this example, it suffices indeed to reduce the mean degree of the network from 〈*k*〉 = 6 to 〈*k*〉 = 3 to switch from local ICW propagation that recruits < 100 astrocytes (*left panel*), to regenerative long-range ICW propagation which activates hundreds of cells (*right panel*).

The dependence of ICW extent of propagation on the network’s mean degree (〈*k*〉, namely on the average number of connections per astrocyte, is further investigated in Figure 3C for all network topologies. It may be appreciated how ICW propagation generally decreases with larger 〈*k*〉: that is, increasing astrocytic connectivity hinders ICW propagation in our networks, independently of their topology. A closer inspection of the figure however allows distinguishing between two classes of networks based on ICW propagation: spatially unconstrained (*black* markers) vs. spatially constrained networks (*red* markers). Here, we dub as “spatially-unconstrained” those networks that can have long-distance connections, but where ICWs activate only few tens of astrocytes. These include for example Erdős-Rényi networks (*black downward triangles*), scale-free networks (*black upward triangles*) or shortcut networks with rewiring probability *p*_*s*_ = 0.3 (*black diamonds*). Conversely, “spatially constrained” networks include link-radius (*red circles*) or regular-degree networks (*red squares*), as well as shortcut networks with *p*_*s*_ = 0 (*red diamonds*), whose connections between astrocytes are locally confined, but where ICWs can recruit > 100 cells. Based on this classification, one may note that the difference between spatially-constrained and spatially-unconstrained networks in terms of the number of cells activated by ICWs can range up to ten folds.

A further useful measure to characterize network connectivity is the network’s mean shortest path *L*. Specifically, this measure can be adopted to quantify the degree of spatial constraining of a network, inasmuch as *L* decreases when short-distance connections are rewired to long-distances ones in networks of the same size (Boccaletti et al., 2006). In this perspective, the differences in ICW propagation shown in Figure 3B can also be correlated with the fact the network in the right panel has a larger value of *L*, and thus contains more short-distance connections than the network in the left panel.

These considerations are further elaborated in Figure 3D where the extent of ICW propagation is shown as a function of the network’s mean shortest path for different astrocytic connectivities. In contrast with what is observed for the mean degree 〈*k*〉 (Figure 3C), the extent of ICW propagation generally increases with *L*. Nonetheless, our distinction between spatially-constrained and spatially-unconstrained networks holds true. It may be noted in fact that, as *L* increases, only link-radius, regular-degree and shortcut networks allow for ICWs that recruit > 100 cells, whereas other network topologies do not. Large *L* values indeed imply dense local, short-distance connections between cells, which can only exists in networks whose topology is subjected to strong spatial constraints.

Overall, the analysis of ICW propagation in our 3d network models predicts that ICW propagation is hindered in astrocytes networks with a large average number of connections per cells and that contain long distance connections (Lallouette et al., 2014). These results are somehow at odds with the notion, supported by studies on neuronal networks models, that small values of mean shortest path and long-distance connections could instead promote signal propagation (Zanette, 2002; Roxin et al., 2004; Dyhrfjeld-Johnsen et al., 2007). This suggests that the principles at play in ICW propagation in astrocyte networks could be different from those involved in action potential propagation in neuronal networks. We focus on these principles in the next section.

### 3.3 Mechanisms of Ca^2+^ wave propagation

At the core of GJC-mediated Ca^2+^ wave propagation is Ca^2+^-induced Ca^2+^ release from the endoplasmic reticulum in the activated astrocytes. This process requires an initial threshold concentration of intracellular IP_3_ to be triggered (Chapter 2). Since in unactivated (resting) astrocytes, endogenous production of IP_3_ by PLCδ is equilibrated by IP_3_ degradation, the only other mechanism that in our model can account for intracellular IP_3_ variations is GJC-mediated IP_3_ diffusion. Hence, only if the inward flux of IP_3_ by diffusion is sufficiently higher than its outward flux, IP_3_ can accumulate in the cytosol of an astrocyte up to the threshold to trigger CICR. When this occurs, the astrocyte gets activated and lies on the front of the ICW.

Consider the cartoon of ICW propagation in Figure 4A, where cells **A, B** and **E** lie on the front of an ICW (*green squares*) that is propagating from left to right through the portion of the depicted network. Because IP_3_ accumulation in these cells must precede their activation, we can equally think of ICW propagation to be driven by the front of intercellular IP_3_ accumulation. In this fashion, what determines if the ICW will propagate to cells **C** or **D** is whether IP_3_ will next accumulate in those cells. With this regard, GJC-mediated diffusion of IP_3_ is such that IP_3_ travels against its gradient. Hence, IP_3_ accumulation in **C** or **D** depends on two diffusive fluxes: (1) a large influx from activated cells **A, B** and **E** (*thick red arrows*) driven by the supposedly larger IP_3_ concentration found in those cells with respect to unactivated cells **C** and **D**; and (2) an outgoing flux to other unactivated cells in the network (*blue arrows*), which grows as intracellular IP_3_ increases in **C** and **D**. In this example, the IP_3_ flux incoming to **C** or **D** ensues from IP_3_ diffusion from only two activated cells (or “IP_3_ sources”), i.e. **A** and **B** for **C**; and **A** and **E** for **D**. Similarly, because both **C** and **D** are connected only to two unactivated neighbors, these latter, akin to “IP_3_ sinks,” control the strength of the IP_3_ flux coming out from **C** and **D**. In general though, it is reasonable to assume that the total inward and outward IP_3_ fluxes of an astrocyte in the network depend on its number of connections, and are thus correlated with the network’s mean degree 〈*k*〉.

**Figure 4:**
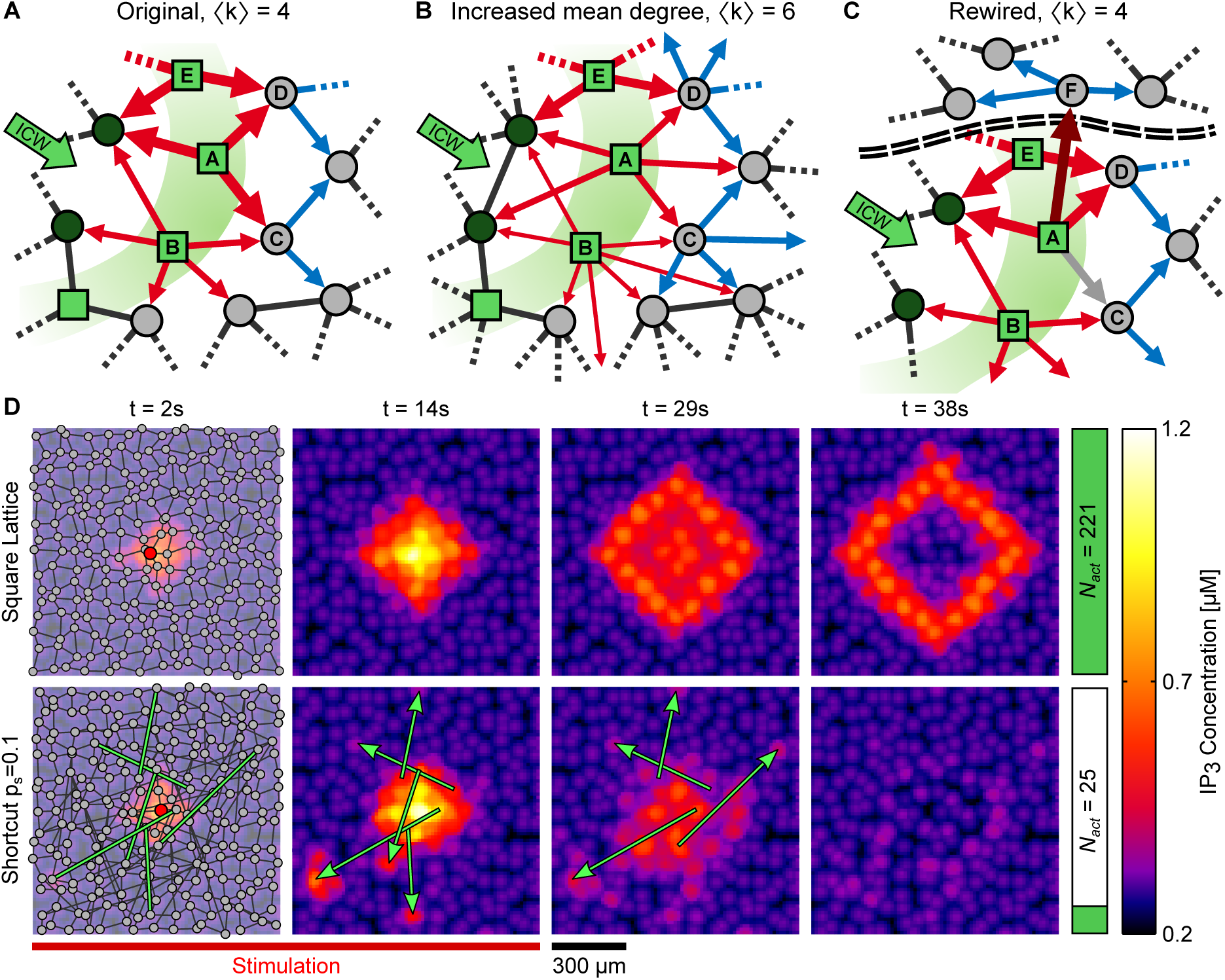
GJC-mediated mechanism of ICW propagation. **A** Propagation of an ICW front (*green squares*) to an astrocyte depends on IP_3_ fluxes mediated by IP_3_ diffusion that comes in (*red arrows*) and out of the cell (*blue arrows*). Unactivated cells (*gray circles*) act as “IP_3_ sinks,” thereby hindering intracellular IP_3_ accumulation. In this fashion, compared with cell A, cell B is less likely to activate other cells for its larger number of connections with unactivated cells. Vice versa, cell D is more likely to get activated than cell C because has fewer unactivated neighbors than this latter. **B** An increase of the network’s mean degree to 〈*k*〉 = 6 introduces additional connections that reduce incoming IP_3_ fluxes to cells C and D, making these cells less likely to get activated (and thus to get recruited by the ICW). **C** Similarly, replacing the local connection between A and C (*gray arrow*) by a long-distance connection between A and F (*dark red* arrow), makes cell C receive less IP_3_, “dumping” the missing IP_3_ flux to an unactivated region of the network far from the wave front (separated by *double dashed lines*). **D** Snapshots of intracellular IP_3_ dynamics at different time instants in two 2d networks shown in the *leftmost panels*). In the shortcut network with *p*_*s*_ = 0.1, the presence of long-distance connections (*green edges and arrows*) makes IP_3_ diffuse away from the wave front (*bottom row*, compare snapshots for *t* = 14s and *t* = 29s). This ultimately results in a considerably lower number of astrocytes activated by an ICW (N_act_, *rightmost vertical bars*) with respect to the square lattice. ICW propagation was triggered stimulating the astrocyte in *red (leftmost panels*) for 0 ≤ *t* ≤ 25 s (*bottom red bar*). Reproduced with permission from Lallouette et al. (2014).

To illustrate this, consider the same cells yet in a network where the mean degree is increased to 〈*k*〉 = 6 (Figure 4B). Astrocytes **A, B** and **E** are now likely weaker sources of IP_3_ for cells **C** and **D**, since there exist additional pathways for IP_3_ diffusion out of them which compete with those from **A, B** and **E** to **C** and **D** (*red arrows*). In turn, **C** and **D** receive less IP_3_ so that they are less likely to reach the IP_3_ threshold for CICR activation. This is also exacerbated by the fact that these cells are somehow larger IP_3_ sinks, insofar as they experience a larger outward flux of IP_3_ for their larger number of unactivated connected neighbors (*blue* arrows).

Similar arguments also hold in the case of a decrease of the network’s mean shortest path **L**. In the previous section we saw how this quantity correlates with the existence of long-distance astrocytic connections. Accordingly, we present in Figure 4C the same network of panel 4A except for rewiring the connection between **A** and **C** (*gray arrow*) by a long-distance connection between **A** and the astrocyte **F** (*dark red arrow*), which we imagine to be in some part of the network far from the ICW front of propagation, and marked by the *dashed double line*. In this scenario, cell **C** is clearly less likely to get activated for the reduced IP_3_ influx that it receives due to the missing connection with cell **A**. The IP_3_ flux from **A** to **F** on the other hand, is also likely not as effective in promoting CICR in cell **F** as it would be in **C**, not only because of a missing contribution to IP_3_ influx in this cell from **B**, but also because cell **F** is in a remote part of the network and, as such, connected to many more unactivated cells than **C**. In other words, it is as if the introduction of the long-range connection between **A** and **F** prevented IP_3_ from accumulating nearby the ICW front, dumping it in a remote, unactivated part of the network.

The interplay between the network’s mean degree and mean shortest path in the regulation of IP_3_ sources and sinks that control ICW propagation, may be promptly elucidated by monitoring intracellular IP_3_ dynamics during ICW propagation (Lallouette et al., 2014). Figure 4D shows snapshots of this dynamics at increasing time instants since stimulation onset (at *t* = 0 in the *red cell*) for two different networks: a strongly spatially-constrained network such as the square lattice with 〈*k*〉 = 4 (*top row*), and a less spatially-constrained network like a shortcut network (*bottom row*), with the same mean degree, yet with ~ 10% of connections being between astrocytes far apart (examples marked in *green*). It may be appreciated how the regular architecture of short-distance connections between cells of the square lattice promotes a compact front of IP_3_ accumulation (*brighter spots* in the *top heat maps*) as the ICW propagates. Conversely, this front is quickly lost for *t* > 14 s in the shortcut network, due to the redistribution of IP_3_ to remote unactivated regions of the network by long-distance connections (*green arrows* in the *bottom heat maps*).

Overall, our modeling of astrocyte networks predicts that the network’s mean degree and mean shortest path could be important determinants of ICW variability of propagation inasmuch as they could control the astrocyte’s propensity to activate and get recruited by an ICW. This propensity ensues from intracellular IP_3_ balance which is regulated by a complex interplay of production, degradation and diffusion fluxes brought forth by activated and unactivated cells. In particular, the number of unactivated neighbors of a given astrocyte could dramatically control its activation as they set the rate of IP_3_ drain from this cell by diffusion. This also accounts for the results discussed in Section 3.1, where we put emphasis on the number of connections with unconnected neighbors as a critical factor to shape the type of Ca^2+^ response of an astrocyte. We can now explain the reason for this result hypothesizing that a cell connected with few unactivated neighbors is likely to accumulate IP_3_ more easily than one with many unactivated neighbors. In this way, that cell not only is likely to get activated faster than the other, but also will display higher frequency of Ca^2+^ oscillations than the cell with many unactivated neighbors, since the frequency of these oscillations grows with intracellular IP_3_ concentration in our model (equations 6–8, but see also De Pittà et al., 2009). Type II vs. type I responses are thus mirrored by these two cells characterized by a different degree of unactivated neighbors.

### 3.4 Comparison of model predictions with experiments

An important prediction of our modeling is that the extent of ICW propagation could be highly variable in astrocyte networks with realistic spatially-constrained topology, like link-radius or regular connectivities, solely depending on the network’s mean degree 〈*k*〉. A 5-fold decrease of 〈*k*〉 from 15 to 3 for example could result in a 100-fold increase in the number of astrocytes recruited by an ICW, from few tens of cells to abot 500 astrocytes (Figure 3). Although no experiments have so far investigated the relationship between ICWs and network topology, there is contingent evidence that astrocytic connectivity could dramatically influence ICW generation and propagation.

Variability of ICW propagation observed in experiments of different astrocyte populations (Charles, 1998; Scemes and Giaume, 2006; Sasaki et al., 2011; Kuga et al., 2011) has indeed been suggested to depend not only on the experimental setup but also on heterogeneities in the connections between astrocytes (Scemes and Giaume, 2006). These heterogeneities have well been characterized for astrocytes in the olfactory bulb which show preferential GJC coupling within rather than outside of glomeruli (Roux et al., 2011). And similar observations have also been made for astrocytes within somatosensory barrels (Houades et al., 2008) and in the stratum pyramidale of the hippocampus (Rouach et al., 2008). In the hippocampus in particular, ICWs could propagate for longer distance in the CA3 region than in the CA1 region (Dani et al., 1992), and it is tempting to speculate that, in light of our modeling, these differences could be due to the fact that CA3 astroytes are known to be less coupled by GJCs (i.e. their (k) is smaller) than their CA1 homologues (D’Ambrosio et al., 1998).

In agreement with the latter hypothesis, is the evidence of reduced ICW propagation in cultures of astrocytoma cells whose coupling was increased by forcing expression of the GJC protein Cx43 (Suadicani et al., 2004). Moreover the fact that ICWs are observed much more frequently in the developing brain (Parri et al., 2001; Weissman et al., 2004; Fiacco and McCarthy, 2006; Scemes and Giaume, 2006; Kunze et al., 2009) rather than in the brain of adult animals (Fiacco and McCarthy, 2006; Scemes and Giaume, 2006) could also be due to developmental differences in GJC expression. Cx30 expression in fact strongly develops between postnatal day 10 (P10) (Aberg et al., 1999) and the third week of life (Rouach et al., 2002). Before this period, neocortical astrocytes are known to be sparsely connected (Aberg et al., 1999) and display ICWs (Iwabuchi et al., 2002). Conversely, during and after this period, the extent of astrocytic ICW propagation seems to drastically reduce. For example, the same stimulation protocol that triggers long-distance ICWs in astrocytes in the CA1 region of the hippocampus before P10, does not after P10, when cell coupling by GJCs is increased (Aberg et al., 1999; Fiacco and McCarthy, 2004).

There is also evidence that expression and permeability of GJC proteins, like Cx30 and Cx43, are regulated by neurons (Rouach et al., 2000; Koulakoff et al., 2008; Roux et al., 2011), possibly by extracellular K^+^ (Pina-Benabou et al., 2001). Remarkably, increases of GJC coupling mediated by extracellular K^+^ were shown to decrease ICW propagation in astrocyte networks (Scemes and Spray, 2012), ultimately suggesting that the (mean) degree of connections of astrocytes in networks is not fixed but rather, depends on local conditions, possibly correlated to ongoing neural activity. Inasmuch as the number of connections of an astrocyte could dictate the characteristics of its Ca^2+^ response to neural activity (Section 3.1), the latter hypothesis opens to the scenario that astrocytes in a network could display different Ca^2+^ responses that depend on activity-dependent modulations of their connectivity. This variegated Ca^2+^ signaling could in turn account for variability of Ca^2+^ activation of individual cells, and ultimately reflect into different modes of recruitment of those cells by ICWs as well as in ICW generation and propagation themselves.

## 4 Simplified modeling of intercellular Ca^2+^ waves

### 4.1 The UAR astrocyte

The biophysical approach considered so far may be effective to model Ca^2+^ signaling and propagation with realistic qualitative features, but has the drawback of limited mathematical tractability for the strong nonlinearity born by the equations of IP_3_-triggered CICR (equations 6–8). Moreover it does not take into account some other aspects of astrocytic Ca^2+^ signaling such as for example its complementary spontaneous (stochastic) generation which may remarkably contribute to ICW nucleation (Skupin et al., 2008, but see also Chapter 4). To fill in this gap, we present in the remaining part of this chapter, a further model of astrocytic ICW propagation that includes stochastic Ca^2+^ activation and is amenable to analytical tractability, while retaining elementary biophysical realism. The results following discussed were originally presented in Lallouette et al. (2014) and Lallouette (2014).

We start from the consideration of “realistic” intracellular IP_3_ and Ca^2+^ dynamics simulated by our biophysical model of equations 6–8. Figure 5A shows IP_3_ and Ca^2+^ traces associated with two ICWs that travel from astrocyte 1 (*top)* to astrocyte 2 (*bottom*) via GJC coupling (equation 5), respectively for 20 < *t* < 40s and 110 < *t* < 120s. One can associate each cell, at any time, with one of three possible states: unactivated (U), activated (A) and refractory (R). In the unactivated state, an astrocyte is at rest, meaning that its intracellular IP_3_ and Ca^2+^ concentrations are either at a low equilibrium, or subjected to some subthreshold dynamics without CICR. Hence, either astrocytes in our example are unactivated before the arrival of each ICW, i.e. for *t* < 20 s and for some time *t* < 110 s.

**Figure 5:**
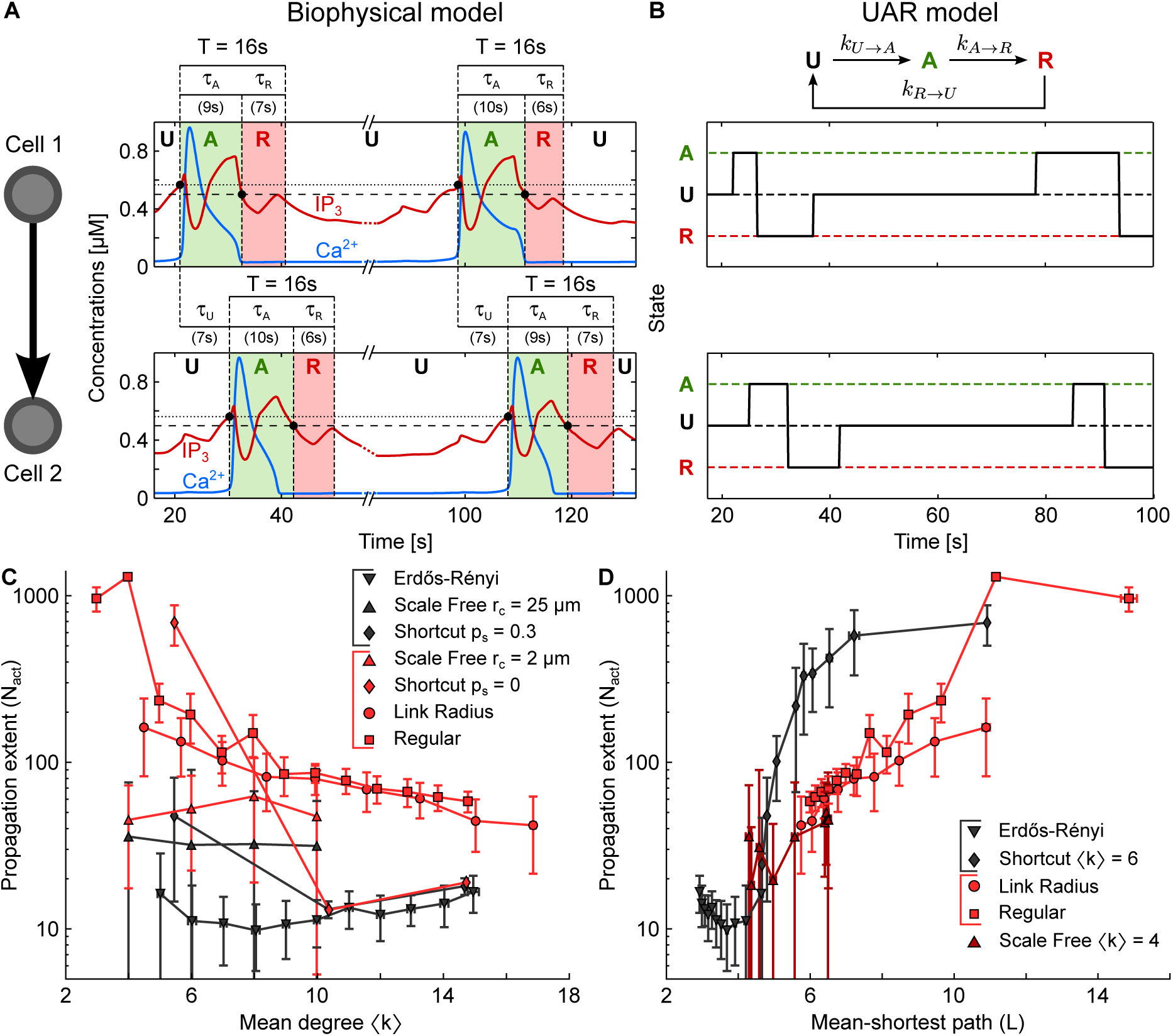
UAR model of ICW propagation. **A, B** Recruitment of an astrocyte by an ICW may be regarded as a three-state process, as exemplified for two connected astrocytes (cell 1, *top row*; cell 2, *bottom row*). Astrocytes are in the unactivated state (U) when at rest. Upon arrival of an ICW, their intracellular IP_3_ (*red traces*) crosses the threshold for CICR initiation (*dotted line*) and cell 1, followed by cell 2, get activated (A, *green-shaded windows*), which is marked by a pulse-like increase of intracellular Ca^2+^ in these two cells (*blue traces*). Following activation, each cell recovers to rest through a refractory period (R, *red-shaded windows*), when their intracellular IP_3_ falls below a supply threshold (*dashed line*). Time constants for each transition may be estimated as following: *τ*_*U*_ coincides with the delay between the Ca^2+^ increases in cell 1 and in cell 2; *τ*_*A*_ is estimated by the time interval from the beginning of the Ca^2+^ elevation to the point where IP_3_ gets below the diffusion threshold; finally, *τ*_*R*_ is derived from *τ*_*A*_ + *τ*_*R*_ = *T*, where *T* = 16 s is the minimum period of Ca^2+^ oscillations in the single astrocyte. Transition rates used in the simulations are obtained averaging over all *τ* values obtained in simulations of the biophysical model in Figure 3. C, D ICW propagation for the same networks of Figure 3 (panels C and D), where astrocytes are modelled instead by the UAR description. The extent of ICW propagation (*N*_*act*_: number of activated astrocytes) generally mirrors qualitative and quantitative characteristics of ICWs simulated in our biophysical network models. Data points±errorbars: mean values±standard deviation over *n* = 20 networks of similar topology. Parameters of the biophysical model and the UAR model are reported in Table C1 and Table C3 respectively. Adapted from Lallouette et al. (2014).

As intracellular IP_3_ crosses the threshold to trigger CICR (*dotted lines*), the astrocytes get activated, displaying a large pulse-like increase of their intracellular Ca^2+^ (*green-shaded windows*). For the previously discussed arguments however (Section 3.3), these cells can stay in this active state of CICR generation as long as IP_3_ supply, by GJC-mediated diffusion from other cells in the network, is large enough to guarantee intracellular accumulation of IP_3_ up to the CICR threshold. This “threshold for IP_3_ supply,” is generally lower than the CICR threshold (*dashed lines*), and can roughly be estimated by the sum of the resting intracellular IP_3_ concentration (~ 0.3 μΜ in our biophysical model) and the gradient ~ 0.2 μΜ for which nearly no GJC-mediated IP_3_ diffusion occurs (see equation 5 and Figure 1C).

Following activation, when IP_3_ drops below the supply threshold, either astrocytes are found in a refractory state (*red-shaded windows*), whereby they do not transmit IP_3_ to other astrocytes but cannot get activated again yet. Finally, as IP_3_ (and Ca^2+^) levels approach their resting values, the cells recover to their unactivated state.

The cycling of an astrocyte through unactivated, activated, refractory and back to unactivated states can be formalized in a simple Markov model, dubbed “UAR model” after the initial of its three states, which is schematized on top of Figure 5B. There, we assume constant rates *k*_*A*→*R*_ and *k*_*R*→*U*_, for the transition of the cell respectively from activated to refractory, and from refractory to unactivated, since these transitions are mainly dictated by the cell’s biophysical properties (Chapter 5). Accordingly, we set *k*_*A*→*R*_ = 1/*τ*̄_*A*_ and *k*_*R*→*U*_ = 1/*τ*̄_*R*_, where *τ*̄_*A*_ is the average time of astrocytic activation during ICW propagation, and *τ*̄_*R*_ = *T* – *τ*̄_*A*_ is the average refractory period, estimated from the minimum period (*T*) of Ca^2+^ oscillations in a single astrocyte.

In contrast, the transition from the unactivated state to the activated one depends on the cell’s intracellular IP_3_ balance which is predominantly altered in resting conditions by IP_3_ diffusion from other cells taking part in ICW propagation. Thus we model the rate of this transition, i.e. *k*_*U*→*A*_, as a function of the states of the cell’s neighboring cells. With this aim, we define the efficacy of the *i*-th astrocyte to supply IP_3_ by

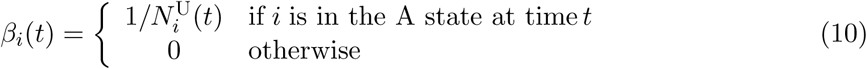

where 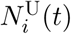 represents the number of unactivated neighbors of *i* at time *t*. The conditional definition of *β*_*i*_(*t*) is motivated by our previous analysis where we noted that only activated cells could effectively supply IP_3_ to unactivated cells that are found next on the pathway of a propagating ICW. The exact functional form *β*_*i*_(*t*) is inspired instead by the observation that the magnitude of IP_3_ supply from an activated astrocyte is inversely proportional to the number of unactivated neighbors, in the assumption of identical neighbors and GJC connections of these latter with cell *i* (see Section 3.3).

Building on our previous analysis, we assume that a given cell i gets activated only if the cumulative IP_3_ supply from its GJC-connected neighbors (𝒩_*i*_ in total) exceeds some threshold for activation ϑ_*i*_. Accordingly, we define its rate of transition from resting to activated as

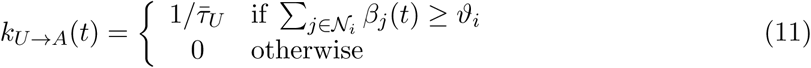

where *τ*̄_*U*_ is estimated from simulations of the biophysical model as the average time needed to activate an astrocyte during ICW propagation, starting from resting intracellular IP_3_ (and Ca^2+^) levels. The threshold *ϑ*_*i*_ can instead be estimated by the minimum number of activated vs. unactivated astrocytes that are connected to a given cell and are required for its activation. In particular it may be shown that this threshold is almost linearly dependent on the cell’s number of connections *k*_*i*_, so that we approximate it here by

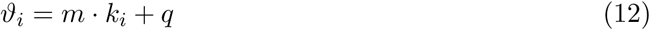

where the parameters *m* and *q* are numerically estimated and depend on the nature of GJC connections (Lallouette et al., 2014).

The UAR model built in the above fashion is reminiscent of SIR models of disease spread (Newman, 2003), and SER models of activity propagation in excitable media (Müller-Linow et al., 2008), yet with an important difference. While in SIR and SER network models, activation of one node of the network generally depends on that node’s immediate neighbor characterized by one degree of separation from the node, in our description, activation of an astrocyte also depends on cells with two degrees of separation from it. For the definition of *β*_*i*_ (equation 10), these cells control in fact the extent at which the connected neighbors of that astrocyte can supply it by enough IP_3_ to trigger its activation.

To verify that the UAR model can reproduce essential predictions provided by our previous biophysical model, we use it to simulate ICW propagation in 3d networks with the same topological features as those considered in Figure 3. The results of these simulations are reported in panels C and D of Figure 5, where it may be seen that the functional dependence of the extent of ICW propagation on the network’s mean degree (〈*k*〉) and mean shortest path (*L*) qualitatively resembles the behavior previously observed for our biophysical model, both in spatially-constrained and spatially-unconstrained networks (cp. panels C and D in Figure 3). The only exception, possibly due to simplifying modeling assumptions on the choice of *β*, is represented by scale-free networks with short-distance connections (*r*_*c*_ = 2 μm, *upward red triangles* in Figure 5B) which allow ICW propagation for > 100 cells for low 〈*k*〉 values, whereas in the biophysical model did not.

### 4.2 Shell propagation model

A close inspection of Figures 3C and 5C reveals a peculiar phenomenon: two network types with very similar spatially-constrained topologies, like cubic lattices obtained from shortcut networks with *p*_*s*_ = 0 (and *l* = 1, see Appendix A), and regular degree networks with 〈*k*〉 = 6, exhibit however a very different behavior, with the former supporting ICWs that could activate 10-fold more cells than in the latter (Figure 6A, *pink* vs. *dark green bars*). It may be hypothesized that, this is due to the fact that cubic lattices with 〈*k*〉 = 6 have a mean shortest path *L* ≈ 11, while regular degree networks with same 〈*k*〉 values associate with *L* ≈ 8.8. Nonetheless, as shown by the *light green bar* of the histogram in Figure 6A, reducing 〈*k*〉 to 4 in these latter networks, so as to obtain *L* values comparable to those in cubic lattices, only marginally increases the extent of ICW propagation in regular-degree networks. This ultimately suggests that the topolgical differences in terms of different 〈*k*〉 and *L* values cannot fully explain variability of ICW propagation and thus other aspects of the network’s architecture must be at play.

**Figure 6:**
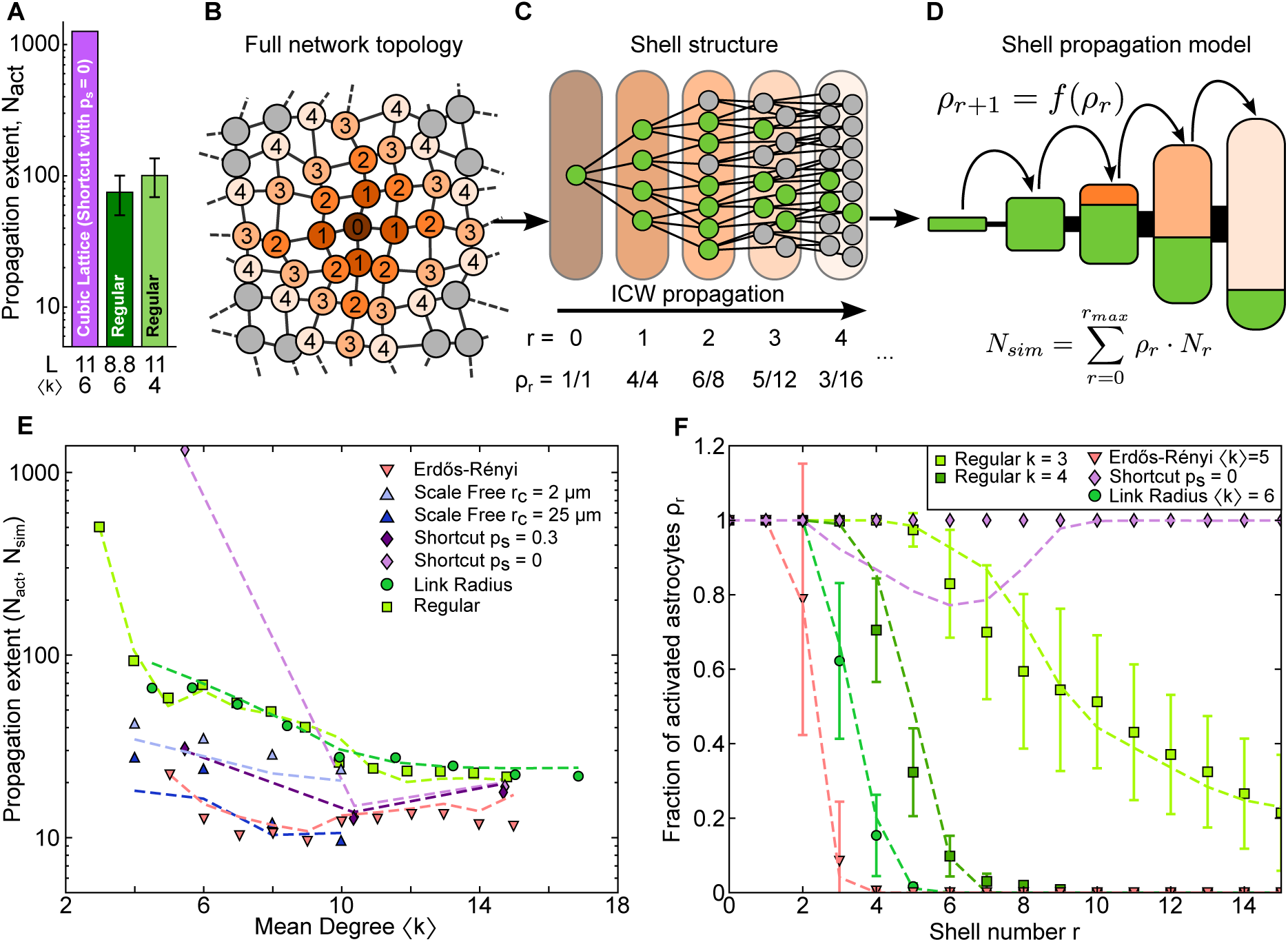
Shell model of ICW propagation. **A** Quantification of ICW propagation by the number of activated cells (*N*_*act*_) in cubic lattices (*pink bar*) vs. regular networks (*green bars*). The 10-fold larger number of activated astrocytes in cubic lattices than in regular networks cannot simply be ascribed to differences in the mean degree (〈*k*〉) and shortest path length (*L*, *bottom values*). Cubic lattices were constructed by shortcut networks with *p*_*s*_ = 0 and *l* = 1 (see Appendix A). **B–D** Shell model of propagation. **B** Sample decomposition of a network neighbor around cell ‘0’ (*umber circle*) by four concentric shells (colors/labels from *brown*/‘1’ to *pale orange*/‘4’). **C** Nodes are grouped by their shell distance r from the reference cell ‘0’. In this fashion, ICW propagation is by activation of astrocytes (*green circles*) from inner to outer shells (i.e. from cell ‘0’ to shell ‘4’ and beyond). **D** At early stages of propagation, the fraction (*ρ*_*r*_) of activated astrocytes per shell is close, if not equal to unity (i.e. all cells are activated – *N*^*r*^ cells in total per shell), but it decreases as the wave propagates through outer shells. For a given shell *r* + 1, this fraction can be recursively expressed as function of the fraction of activated cells in the inner shell *r* (*top equation*). The total number of activated cells by shell propagation of an ICW is quantified by *N*_*sim*_ (*bottom formulaa*). **E** The number of activated cells (*N*_*sim*_) and F the fraction of activated cells (*ρ*_*r*_) estimated by the shell model (*dashed lines*) are superimposed on values from simulations of biophysical networks (*data points* from Figure 3C). The shell model of propagation well predicts ICW extent simulated by biophysical modeling. Model parameters as in Table C3.

Both cubic lattices and regular-degree networks were constructed in a similar way: in the former, astrocytes first were linked to their nearest neighbors and their positions were then jittered; in the latter, the order of these operations was reversed. Thus differences in their architectures are subtle and likely relate to the details of local connections of individual cells with their neighbors. To describe these differences, we introduce the notion of propagation shell. Specifically, we define the *r*-th shell with respect to a reference astrocyte, as the ensemble of cells whose topological distance from that astrocyte is *r*.

Figure 6B shows the first four shells of a reference *umber* astrocyte (labeled by ‘0’) in a square lattice. The first shell (*brown* cells with ‘1’ label) is made of the astrocytes that are directly connected with astrocyte ‘0’. The second shell (*orange* cells with label ‘2’) is composed instead by all the cells with two degrees of separation from the reference astrocyte, that is the shortest path to go from them to astrocyte ‘0’ is 2; the third shell (*light orange*) is made of all astrocytes with three degrees of separation from cell ‘0’ and so on.

When stimulating astrocyte ‘0’, the ICW that generates from this cell and propagates to its periphery, may be thought as the result of the progressive activation of shells 1, 2 and so on. Figure 6C illustrates this concept, showing in *green* the fraction of astrocytes per shell that get activated by the ICW. It may be noted from Figure 6D that the cells in the first two shells are nearly all activated but, as the wave propagates to outer shells (i.e. *r* > 2), the ratio of activated cells per shell (*ρ*_*r*_) quickly drops, ultimately halting ICW propagation beyond the fourth shell.

By the same arguments previously exposed in Section 3.3, what determines the extent of activation of a shell, and whether an ICW propagates to the next one, is respectively the IP_3_ supply to and from that shell. This supply can be estimated by means of the UAR model, resolving for the network’s shell structure, including the number of cells per shell (*N*_*r*_), and the fraction of activated astrocytes per shell (*ρ*_*r*_) against the number of unactivated cells therein (*N̂*_*r*_). Denoting by 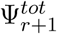 the average total IP_3_ supply to an astrocyte in shell *r* + 1 from shell *r*, this quantity may be regarded as the sum of two terms in general: (i) an endogenous IP_3_ supply by IP_3_ production and diffusion from shell *r*, 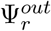; and (ii) an exogenous IP_3_ supply to shell *r* + 1 directly due to the stimulation protocol, i.e. 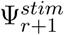 – clearly, the farther the shell is from the stimulated cell, the lesser is the IP_3_ directly supplied to it by the applied stimulus. That is

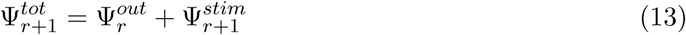

Then, the IP_3_ supplied by the *r*-th shell can be thought to be proportional to the ratio between the number of activated astrocytes (or IP_3_ sources) in that shell and the number of unactivated cells (or IP_3_ sinks) in the proximal shell *r* + 1, i.e.

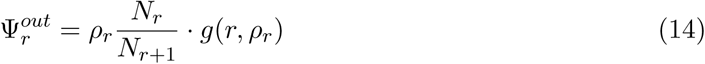

where the multiplicative factor *g*(*r*, *ρ*_*r*_) accounts for the size of shells *r* – 1 and *r* + 1 and the connections of cells therein, with themselves and with cells in other shells. Finally, based on the previous equations, the fraction of activated cells in shell *r* + 1 is given by

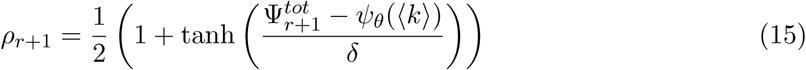

where *ψ*_*θ*_(〈*k*〉) is homologous of the activation threshold *ϑ* in equation 11 for the *r*-th shell. The fraction of activated astrocytes per shell is thus a sigmoid function of IP_3_ supplied to that shell by inner shells. It approaches 1 when this IP_3_ exceeds the threshold *ψ*_*θ*_(〈*k*〉), which corresponds to the ideal scenario of all cells in the shell being activated, and to an ICW that propagates to shell *r* + 1 in a perfectly regenerating fashion. Conversely, it tends to 0 when 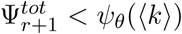, ensuing in partial ICW propagation to shell *r* + 1, which possibly preludes to wave death in the following shells. The slope of the transition between partial (vanishing) ICW propagation (i.e. *ρ*_*r*+1_ → 0) and fully regenerative ICW propagation (*ρ*_*r*+1_ → 1) is controlled by the parameter *δ*. The detailed derivation of equations 13–15 may be found in the online Supplementary Text (Appendix ??).

Using equation 15, we can recursively compute the activation ratio of concentric shells of ICW propagation, ultimately estimating the extent of ICW propagation without the need to simulate whole networks. Figure 6E reports the results of this estimation, where the extent of ICW propagation is quantified by the number of astrocytes expected to get activated by an ICW, i.e. *N*_*sim*_ = Σ_*r*_ *N*_*r*_*ρ*_*r*_. Comparison of *N*_*sim*_ values obtained by our shell description of ICW propagation (*dashed lines*) with those for the number of activated astrocytes from simulations of the biophysical model (*data points* from Figure 6C), reveals a close correspondence of our analytical estimation with simulations. Further analysis (Figure 6F) also shows that the estimated fraction of activated astrocytes per shell is in good agreement with the majority of data from ICW propagation simulated in biophysical network models.

Overall, the analysis presented in this section puts emphasis on the importance of regional features of cell connectivity mirrored by the shell structure of astrocyte networks, as a crucial factor in shaping ICW propagation. This propagation appears to depend on the shell-to-shell GJC-mediated diffusion IP_3_ by equation 15 in a strongly nonlinear fashion, with inner shells driving activation of outer shells during ICW radial propagation towards the network periphery of a stimulated cell. In this fashion, as far as the activation of inner shells is guaranteed, an ICW could regenerate and propagate across large portions of the network, if not the whole network. On the other hand, a simple change of the stimulus protocol, resulting in an alteration of IP_3_ supply that can no longer robustly activate astrocyte shells that are proximal to the stimulus site, would cause ICWs to propagate only for short distances. This could ultimately explain why the same astrocyte networks some times display local Ca^2+^ activity, spatially confined to ensembles of few activated cells, (Sasaki et al., 2011) and some other times long distance ICWs, engulfing hundreds of cells (Kuga et al., 2011).

## 5 Conclusions

The computational arguments presented in this chapter pinpoint to topological determinants of signal propagation in astrocyte networks that substantially differ from those at play in neural networks. While neurons communicate by electrical signals using distinct pools of neurotransmitters, found at each of their synapses, astrocytes propagate Ca^2+^ signals by a complex exchange of IP_3_ fluxes, controlled by the exact spatial arrangement of IP_3_ sources and sinks ensuing from activated vs. unactivated cells. Hence, while increasing the number of connected neighbors in a neural networks would be tantamount, in our description, to add new synapses thereby increasing cell excitability; in an astrocyte network instead, this could reduce IP_3_ supply to individual cells, hindering cell activation and ICW propagation.

In agreement with this view, increasing the average number of connections per cell (i.e. the network’s mean degree) in models of networks of excitatory neurons was suggested to promote neural synchronization, that is the coherent activation of neuronal ensembles (Wang et al., 1995; Golomb and Hansel, 2000; Olmi et al., 2010; Luccioli et al., 2012; Tattini et al., 2012). In contrast, we showed how cell hubs (that is, cells with a large number of connections), and long-distance connections between astrocytes could dramatically break ICW fronts, substantially reducing the extent of their propagation. These differences in terms of topology vs. dynamics in neural vs. astrocyte networks brought forth in this chapter, contribute to shed new light on functional and organizational principles beyond astrocyte networks whose topological features are notoriously very different from those on neural networks (Bushong et al., 2002).

## Appendix A Simulations of 3d astrocytic networks

### A.1 Construction of networks with different topology

The five different topologies for 3d astrocytic networks considered in this chapter were constructed as following detailed (see also Lallouette et al., 2014).

- *Link radius networks* were constructed connecting each astrocyte *i* to all cells contained in a sphere of radius *d* centered on *i*. The degree distribution of these networks displays some variability around the mean degree 〈*k*〉, due to preliminary jitter of astrocyte locations in the absence of highly connected cells.
- *Regular degree networks* were developed connecting each astrocyte to its *k* nearest neighbors while forbidding links longer than *d*_*max*_ = 150 μm. In doing so, connections were established in *k*_*reg*_ iterations to avoid directional biases. Namely, all nodes were randomly taken once per iteration *m* and linked to the nearest node *i* with degree *k*_*i*_ < *m* ≤ *k*_*reg*_ and located within *d*_*max*_ from the selected node.
- *Shortcut networks* were constructed in a way similar to small-world networks (Watts, 1999). We started by positioning astrocytes on a cubic lattice with internode distance *a*, linking each cell to its nearest neighbors at distances that were multiples of *a* up to *l* times. We then rewired each connection with probability *p*_*s*_ thereby randomly assigning one of its endpoint. Finally, we jittered the nodes positions as explained in the main text.
- *Spatial scale-free networks* were incrementally built by spatially-constrained preferential attachment (Barthélemy, 2010). Briefly, astrocytes were progressively included in the network, one by one, and connected with *m*_*sf*_ cells. Each connection between a new astrocyte *i* and a target cell *j* was established with probability *p*_*i*→*j*_ ∝ *k*_*j*_ exp(–*d*_*ij*_/*r*_*c*_); where *k*_*j*_ is the degree of the target cell *j*, *d*_*ij*_ represents the Euclidean distance between astrocytes *i* and *j*, and *r*_*c*_ sets the range of interaction between cells in the space. Small values of *r*_*c*_ result in connections between astrocytes that are limited to their neighbors, while large rc values allow establishing long distance connections. Spatially-constrained preferential attachment may also produce some highly-connected astrocytes or ‘hubs’.
- *Erdős-Rényi networks* were built by linking each astrocyte pair with probability *p*, independently of their distance and existing degree. These networks are the only ones in our analysis that do not bear any spatial constraint.

Depending on whether *r*_*c*_ (respectively *p*_*s*_) is large or not, spatial scale-free networks (respectively shortcut networks) can be regarded either as spatially-constrained networks or as spatially-unconstrained networks. Due to random wiring, some of the above procedures could result in disconnected networks. To minimize this scenario, we iterated the wiring procedure to ensure that, in our networks, disconnected node pairs accounted for < 2% of all possible node pairs. Parameters used to build the different networks in the simulations discussed in Section 3.2 are detailed in Table C2.

### A.2 Numerical procedures

Each network model consisted of 3*N* = 3993 ODEs which we numerically solved by 4^th^ order Runge-Kutta integration with a time step of 0.01 s. The extent of ICW propagation (*N*_act_) was quantified by the number of astrocytes that were activated at least once, where an astrocyte was considered to be activated whenever its Ca^2+^ concentration exceeded 0.7 μM. Each network model was produced into *n* = 20 different realizations, and mean degree (〈*k*〉) and mean shortest path length (L) of each network model were averaged over realizations.

To generate ICWs, we stimulated the cell whose location was the closest to, if not coincident with the center of the 3d cubic lattice containing the network. Stimulation was delivered for 0 ≤ *t* ≤ 200 s connecting an IP_3_ reservoir of 2 μM to the central cell and allowing IP_3_ diffusion into that cell according to equation 5.

### A.3 UAR model simulations

In networks with UAR astrocytes, we considered step increases of time by Δ*t* = 0.1 s, simulating a transition from a state *x* to a state *y* (with rate *k*_*x*→*y*_ and *x*,*y* = U, A, R), every time that a random number *r* drawn from a uniform distribution in [0,1] at each Δ*t* was such that *r* ≤ *k*_*x*→*y*_ · Δ*t* In those networks, stimulation of the central cell was deployed forcing activation of its connected neighbors, since this was observed to be case in the majority of networks with biophysically-modeled astrocytes.

## Appendix B Supplementary online material and software

Detailed derivation of the shell model (Section 4.2) can be downloaded from https://github.com/mdepitta/comp-glia-book. The file `Shell.derivation.pdf` is provided along with the original LATEX files within the `doc` folder associated with this chapter. In the same folder the WxMaxima file `ODEsystem.wxmx` is also provided. This file was used to analytically solve the ODE system at the core of the derivation of the shell model (equations 1–3 in the supplementary online text).

Within the same repository the code used for simulations of astrocyte networks presented in this chapter is also provided. The core source code is implemented in C++ and is located in `src` folder. This code must preliminarly be compiled by `make` from this directory. The Python script, `RunSimulations.py` relies on the compiled source code to generate all data sets to plot the figures of this chapter. Depending on the hardware configuration, it might take up to a day to complete all the simulations involved. By default, the software will attempt using all available cores on the local machine. Individual figures can be generated by `Figure_3.py` for Figure 3C and D; `Figure_5.py` for Figure 5C and D; and `Figure_6.py` for Figure 6E and F.

## Appendix C Model parameters used in simulations

**Table C1:** Parameters of the biophysical model.

| Symbol | Description | Value |  | Units |
| --- | --- | --- | --- | --- |
|  |  | GChI | ChI |  |
| *IP<sub>3</sub>R kinetics* |  |  |  |  |
| $d_1$ | IP<sub>3</sub> binding affinity | 0.13 | | μM |
| $O_2$ | Inactivating Ca<sup>2+</sup> binding rate | 0.62 | 0.2 | μM<sup>-1</sup>s<sup>-1</sup> |
| $d_2$ | Inactivating Ca<sup>2+</sup> binding affinity | 1.049 | | μM |
| $d_3$ | IP<sub>3</sub> binding affinity (with Ca<sup>2+</sup> inactivation) | 0.9434 | | μM |
| $d_5$ | Activating Ca<sup>2+</sup> binding affinity | 0.08234 | | μM |
| *Calcium fluxes* |  |  |  |  |
| $C_T$ | Total ER Ca<sup>2+</sup> content | 2 | | μM |
| $\rho_A$ | ER-to-cytoplasm volume ratio | 0.185 | | — |
| $\Omega_C$ | Maximal Ca<sup>2+</sup> release rate by IP<sub>3</sub>Rs | 18.56 | 6 | s<sup>-1</sup> |
| $\Omega_L$ | Maximal Ca<sup>2+</sup> leak rate | 0.3416 | 0.11 | s<sup>-1</sup> |
| $O_P$ | Maximal Ca<sup>2+</sup> uptake rate | 2.7846 | 0.9 | μMs<sup>-1</sup> |
| $K_P$ | Ca<sup>2+</sup> affinity of SERCA pumps | 0.05 | | μM |
| *IP<sub>3</sub> production* |  |  |  |  |
| $O_\delta$ | Maximal rate of IP<sub>3</sub> production by PLCδ | 0.4641 | 0.7 | μMs<sup>-1</sup> |
| $K_\delta$ | Ca<sup>2+</sup> affinity of PLCδ | 0.1 | | μM |
| $\kappa_\delta$ | Inhibiting IP<sub>3</sub> affinity of PLCδ | 1.5 | | μM |
| $O_\beta$ | Maximal rate of IP<sub>3</sub> production by PLCβ | 1.105 | — | μMs<sup>-1</sup> |
| $K_G$ | Glutamate affinity of the receptor | 1.3 | — | μM |
| $K_L$ | Ca<sup>2+</sup>/PKC-dependent inhibition factor | 10 | — | μM |
| $K_{KC}$ | Ca<sup>2+</sup> affinity of PKC | 0.6 | — | μM |
| *IP<sub>3</sub> degradation* |  |  |  |  |
| $\Omega_{5P}$ | Maximal rate of IP<sub>3</sub> degradation by IP-5P | 0.793 | 0.21 | s<sup>-1</sup> |
| $O_{3K}$ | Maximal rate of IP<sub>3</sub> degradation by IP<sub>3</sub>-3K | 13.923 | 4.5 | μMs<sup>-1</sup> |
| $K_D$ | Ca<sup>2+</sup> affinity of IP<sub>3</sub>-3K | 1 | | μM |
| $K_{3K}$ | IP<sub>3</sub> affinity of IP<sub>3</sub>-3K | 0.7 | | μM |
| *IP<sub>3</sub> diffusion* |  |  |  |  |
| $F$ | GJC IP<sub>3</sub> permeability | 3.64 | 2 | μMs<sup>-1</sup> |
| $I_\theta$ | Threshold IP<sub>3</sub> gradient for diffusion | 0.15 | 0.3 | μM |
| $\omega_I$ | Scaling factor of diffusion | 0.05 | | μM |
| *Synaptic glutamate release* |  |  |  |  |
| $\Omega_f$ | Rate of synaptic facilitation | 2 | — | s<sup>-1</sup> |
| $\Omega_d$ | Rate of recovery of released synaptic vesicles | 1 | — | s<sup>-1</sup> |
| $\Omega_G$ | Glutamate clearance rate | 60 | — | s<sup>-1</sup> |
| $U_0$ | Basal probability of synaptic glutamate release | 0.25 | — | — |
| $\rho_C$ | Ratio of synaptic vesicles and mixing volumes | $6.5 \times 10^{-4}$ | — | — |
| $G_T$ | Glutamate content of readily releasable vesicles | 200 | — | mM |
| $\rho_{so}$ | Glutamate spill over fraction to the astrocyte | 0.075 | — | — |

**Table C2:**
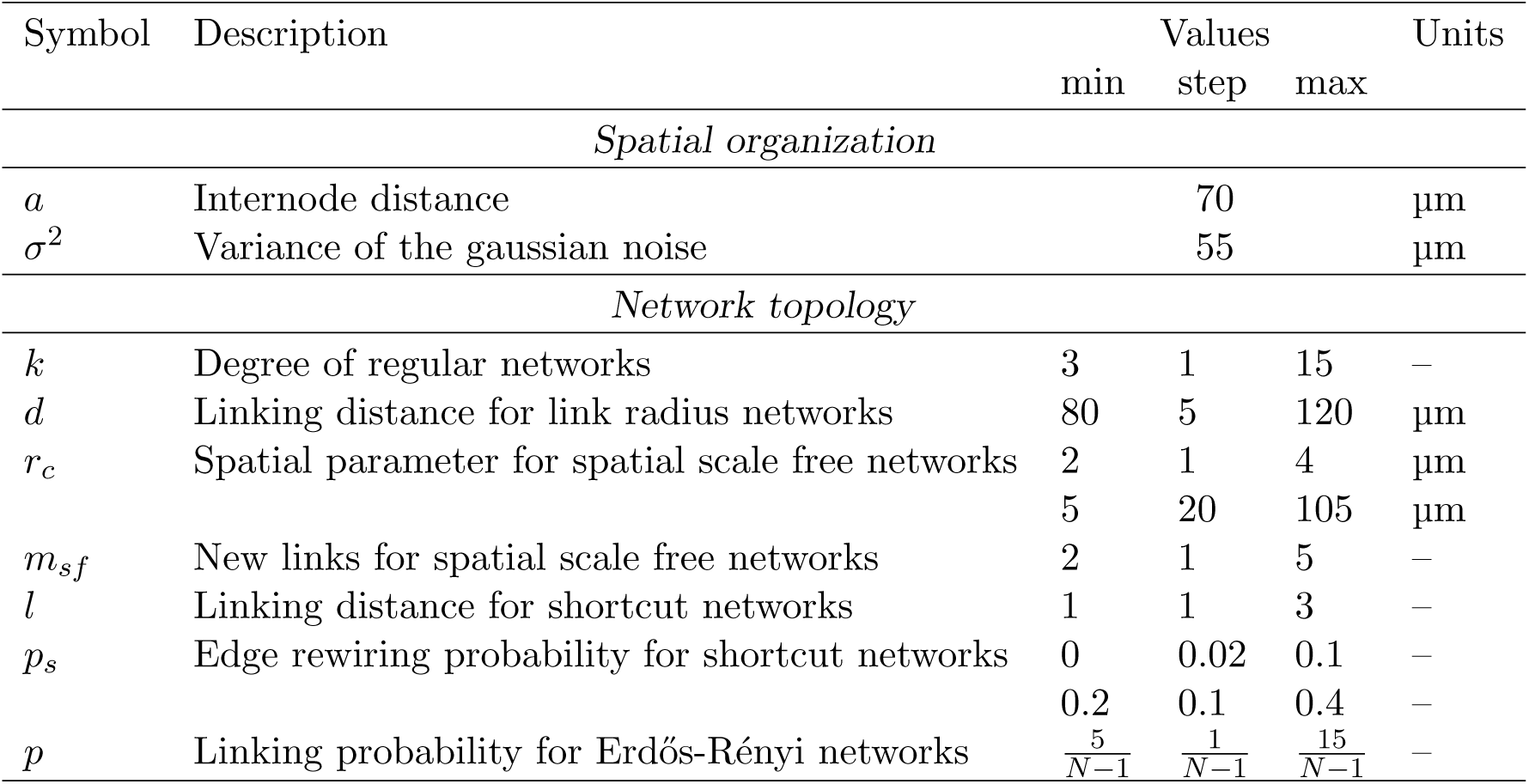
Spatial and topological parameters of the astrocyte network model.

| Symbol | Description | Values |  |  | Units |
| --- | --- | --- | --- | --- | --- |
|  |  | min | step | max |  |
| *Spatial organization* |  |  |  |  |  |
| $a$ | Internode distance | | 70 | | $\mu\text{m}$ |
| $\sigma^2$ | Variance of the gaussian noise | | 55 | | $\mu\text{m}$ |
| *Network topology* |  |  |  |  |  |
| $k$ | Degree of regular networks | 3 | 1 | 15 | — |
| $d$ | Linking distance for link radius networks | 80 | 5 | 120 | $\mu\text{m}$ |
| $r_c$ | Spatial parameter for spatial scale free networks | 2 | 1 | 4 | $\mu\text{m}$ |
| | | 5 | 20 | 105 | $\mu\text{m}$ |
| $m_{sf}$ | New links for spatial scale free networks | 2 | 1 | 5 | — |
| $l$ | Linking distance for shortcut networks | 1 | 1 | 3 | — |
| $p_s$ | Edge rewiring probability for shortcut networks | 0 | 0.02 | 0.1 | — |
|  |  | 0.2 | 0.1 | 0.4 | — |
| $p$ | Linking probability for Erdős-Rényi networks | $\frac{5}{N-1}$ | $\frac{1}{N-1}$ | $\frac{15}{N-1}$ | — |

**Table C3:**
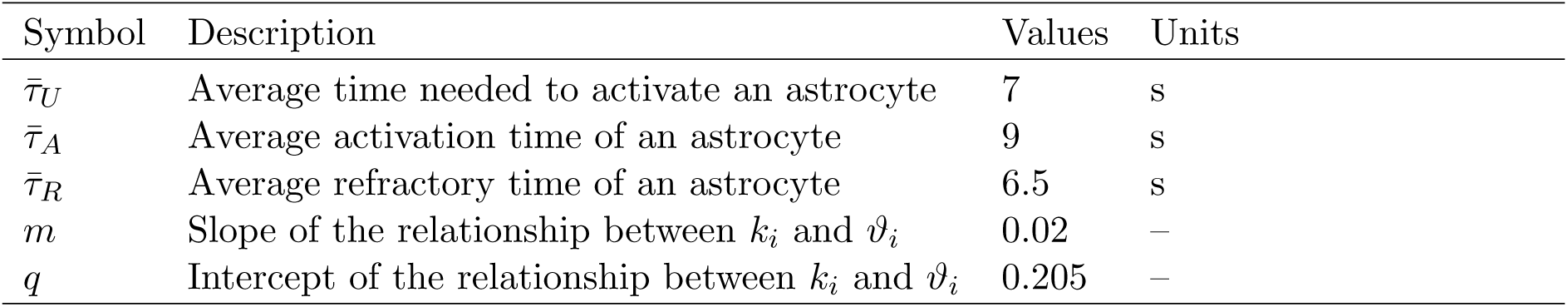
UAR model parameters.

| Symbol | Description | Values | Units |
| --- | --- | --- | --- |
| $\bar{\tau}_U$ | Average time needed to activate an astrocyte | 7 | s |
| $\bar{\tau}_A$ | Average activation time of an astrocyte | 9 | s |
| $\bar{\tau}_R$ | Average refractory time of an astrocyte | 6.5 | s |
| $m$ | Slope of the relationship between $k_i$ and $\vartheta_i$ | 0.02 | – |
| $q$ | Intercept of the relationship between $k_i$ and $\vartheta_i$ | 0.205 | – |

**Table C4:** Shell model parameters.

| Symbol | Description | Values | Units |
| --- | --- | --- | --- |
| $\frac{\Omega_I}{F}$ | Effective degradation rate | 2.9883 | – |
| $S$ | IP<sub>3</sub> supplied to the stimulated cell | 2169.32 | – |
| $\eta$ | Power law exponent | 2.62 | – |
| $A$ | Slope of $\psi_\theta(\langle k \rangle)$ | $2.622 \cdot 10^{-15}$ | – |
| $B$ | Intercept of $\psi_\theta(\langle k \rangle)$ | 0.35061 | – |
| $\delta$ | Slope of the shell activation function | 0.05388 | – |

